# Astroglial Dysfunction in Models of CDKL5 Deficiency Disorder

**DOI:** 10.64898/2026.06.09.730164

**Authors:** Ceri Pickering, Adriana Bakoulina, Faye McLeod, Angeliki Pantziarou, Zhi Yi Alicia Seet, Alishba Saleemi, Yimiao Wu, Gavin J Clowry, Christopher J A Cowie, Maria Kinali, Nicholas D Mazarakis

## Abstract

CDKL5 Deficiency Disorder (CDD) is a rare developmental epileptic encephalopathy typically caused by loss of function variants in the gene encoding the X-linked serine-threonine kinase CDKL5. CDKL5 is highly expressed in the brain during development, and key neuronal functions of the kinase include cytoskeletal organisation and synaptic stability. However, at present, little is known about the function of astroglia in CDD. Given the importance of these cells in synaptic development and homeostasis, as well as dysfunction in other epileptic diseases, it was hypothesised that astrocytes may contribute to CDD pathology. Induced pluripotent stem cells harbouring a CDKL5 loss-of-function mutation (and isogenic controls) were derived from CDD patient fibroblasts and differentiated into astrocytes (iAstros). Analysis of iAstros revealed transcriptomic, proteomic and functional dysregulation in CDKL5-mutant iAstros relating to water transport and immunological function, including a diminished response to TNFα stimulation. Moreover, iAstros showed increased branching and reduced phosphorylation of the known CDKL5 target end-binding protein 2 (EB2) - indicative of disrupted cytoskeletal regulation in a manner similar to CDKL5-null neurons. Finally, we report the generation of novel *in vitro* models of CDD. CDKL5 was knocked down in adult and foetal human organotypic brain slices through transduction with an AAV encoding a novel CDKL5 shRNA. Slices transduced with the CDKL5 shRNA displayed increased spontaneous network activity, demonstrating the functionality of this model. Importantly, interrogation of these models revealed dysregulation of key astrocytic proteins congruous with the human glial stem cell model. Consequently, this study describes the generation of novel human models of CDD and their associated astrocytic dysfunction – paving the way for novel discovery and therapeutic intervention.

## Introduction

CDKL5 Deficiency Disorder (CDD) is a rare X-linked genetic disorder; a haploinsufficiency with a frequency of approximately 1 in 40000-60000 newborns [1]. The disease is characterised by intractable early-onset epilepsy, severe motor and cognitive developmental delay, and autism-like behaviours [1, 2]. Seizure type, frequency, and severity vary over time, and may re-emerge differently following treatment with anti-seizure medication, to which CDD patients are highly refractory [3, 4]. Motor and cognitive developmental milestones are severely delayed in CDD, with less than a quarter of patients attaining independent ambulation [1], only 5% able to feed independently, and most patients achieving an IQ score of no more than 40 [5]. Sleep disorders are also common in CDD, with symptoms including extended periods of insomnia, regular night-waking, excessive drowsiness, and sleep breathing disorders [6]. Gastrointestinal and feeding disorders including constipation, acid reflux, and dysphagia are also frequently observed, with a fifth of patients requiring gastrostomy insertions for feeding [1]. Heterozygous female CDD patients exhibit a wide spectrum of phenotypes, from milder forms that allow for independent walking and drug-controlled seizures, to severe forms with intractable seizures, microcephaly, and minimal independent motor control [1]. Males with CDD are four times rarer than female patients and typically present with a more severe phenotype across all groups of symptoms [7]. There is currently no cure for CDD, with available treatments offering modest and largely symptomatic therapeutic effects. Anti-seizure medications are a first-line treatment, with over 70% of patients being on multiple drug therapy [1, 3, 4]. Ganaxolone (Ztalmy®), a positive allosteric GABAA receptor modulator, is the first FDA-approved CDD-specific therapeutic and has been shown to reduce seizure burden in children aged 2 and above by about 30%, is currently in phase III trials [8].

CDD arises due to pathogenic variants in the *CDKL5* gene that result in an absent, non-functional, or poorly functioning CDKL5 protein. The human *CDKL5* gene is located at the Xp22.13 position on the short arm of X chromosome [9], and comprises 27 exons that form 5 different transcriptional isoforms, of which 4 are expressed in the brain [10]. *CDKL5* pathogenic variants are *de novo* and can be classified into four main groups: complete absence of functional protein, missense variants located within the kinase domain, protein-truncating variants (PTVs) located between amino acids 172-781, and PTVs between amino acids 781-905 respectively [11]. The different types are broadly correlated to achievement of developmental milestones and seizure frequency, with complete absence of protein having the most severe outcome [11].

The CDKL5 protein is a serine-threonine kinase that is highly expressed in the central nervous system, particularly in the cerebral cortex, cerebellum, brainstem, thalamus, and hippocampus [12]. It is most highly and widely expressed in neurons in the CNS but is also found in several other CNS cell types and in peripheral tissues [10]. Expression is developmentally regulated and peaks in early postnatal life [13], when synaptogenesis and neuronal maturation occur. Onset of CDD symptoms also typically begin in this period [1]. The protein has a catalytic N-terminal, which contains its ATP-binding site, serine-threonine kinase active site, and TEY motif [14, 15]. It also has a regulatory C-terminal, which contains its nuclear localisation and export signals, enhances the catalytic activity of the N-terminal and is crucial for CDKL5 subcellular localisation [12, 14].

The CDKL5 protein performs key roles in cell proliferation, neuronal migration and development, synaptic function, axon growth, and dendritic morphogenesis [14]. Absence or deficiency of the protein has drastic impacts on the morphology and development of neurons, damaging neuronal circuit connections, and reducing or causing dysmorphic dendritic and axonal architecture [16–18]. As a kinase, the key function of CDKL5 is to regulate downstream cellular process through the phosphorylation of specific targets [15, 19, 20]. Crucial findings from previous studies include the identification of several cytoskeleton-related targets and a growing agreement that targets of CDKL5 should possess the RPXS/T amino acid consensus motif to be phosphorylated by CDKL5.

To date, there have been several attempts to generate effective models of CDD. Primarily, efforts have been focused on mouse models and the differentiation of patient cells *in vitro*, although other models such as the use of rats and zebrafish have also been described in the literature [21–23]. Several CDKL5 knockout mouse models have been generated, that replicate symptoms such as learning and memory impairment, social and motor deficits, autistic-like behaviours, and signalling pathway dysfunction [16, 24–26]. A key caveat with the mouse models of this disorder is the general lack of early spontaneous seizures, whilst the majority of human *in vitro* models lack network connectivity. Nonetheless, models of CDD have facilitated the study of key mechanisms involved in the development of epilepsy and learning and memory deficits, and have aided in the development of potential treatment strategies [27, 28].

Astrocytes play key roles in synapse formation, neuronal signalling, tissue homeostasis, and regulation and stabilisation of the blood-brain barrier (BBB) [29–31]. Glial cells are thought to comprise a 1:1 ratio with neurons, with approximately 20-40% of these being astrocytes [32]. Importantly, astrocytes are responsible for regulating extracellular levels of neurotransmitters such as glutamate and gamma-aminobutyric acid (GABA), which are the primary excitatory and inhibitory neurotransmitters of the CNS respectively, and are highly implicated in epilepsy. Astrocytic dysfunction is a hallmark of the epileptic focus, both in human and animal models of disease [33]. In epileptic tissue, astrocytes lose regular domain organisation [34], become uncoupled [35], and undergo reactive astrogliosis, bearing subsequent significant changes in morphology, number, function, and transcriptional regulation [33, 36].

As CDKL5 expression is predominantly neuronal its possible function in other cell types remains unclear. In our study we set out to generate several models through which to study the effect of CDKL5 deficiency in astrocytes *in vitro*. Firstly, iPSC-derived astrocytes were generated, to understand CDKL5 function in an isolated human cell-type model. Next, to understand whether human and mouse models of CDD astrocytes are comparable, transcriptomic studies were repeated using isolated cortical and hippocampal astrocytes from a mouse model of CDD. Interestingly, analysis of these models and data in the literature seemed to suggest that mouse astrocytes may express a lower basal level of CDKL5 than human astrocytes and therefore are less relevant in CDKL5-null mice. Combined with the fact that these mice do not exhibit spontaneous seizures, the need for better human models of the disorder was therefore highlighted. Whilst iPSC-derived astrocytes as human models of CDD confer some utility, an understanding of how astrocytes may contribute to CDD in a representative network of human CNS cell types could not be inferred due to the single cell-type nature of this model. Therefore, we describe proof-of-concept of a novel model of CDD using foetal and adult human organotypic brain slice cultures. These models may be utilised by future researchers to interrogate the dysfunction of many CNS cell-types at developmental and post-developmental timepoints in CDD, as well as to screen potential therapeutic candidates. The three models presented in this study represent three interdependent studies into CDKL5-deficient astrocytes, each with a distinct value and each suggesting that astrocytes may play a thus-far uncharacterised role in CDD.

## Methods

### Cell lines

All cell culture was performed under sterile conditions. Cells were passaged when approaching confluency and media was replaced every 2-3 days. Human Embryonic Kidney 293T (HEK293T/17 CRL-11268) cells (ATCC) were maintained in DMEM-F12 High Glucose supplemented with 10 % Fetal Bovine Serum (FBS), 2 mM L-Glutamine and 1 X Penicillin/Streptomycin. Normal Human Astrocytes (Lonza CC-2565) were maintained in ABM Astrocyte Basal Medium (Lonza, CC-3186). When cells reached 70-90% confluency, they were dissociated with Trypsin-EDTA (Sigma-Aldrich-Aldrich, T4174) and plated at a density of 5 x 10^3^ cells/cm^2^. C57BL/6 Mouse Astrocytes (ScienCell, M1800-57) were grown at a density of 5000 cells/cm^2^ into tissue culture-treated plasticware coated with 2 µg/cm^2^ poly-L-lysine (ScienCell, 0403). Mouse astrocytes were maintained in ScienCell Astrocyte Medium-animal (ScienCell, 1831) supplemented with 10 % FBS, 1X Astrocyte Growth Supplement-animal (ScienCell, 1882) and 1 X Penicillin/streptomycin, as per manufacturers instructions.

### iPSC lines and astrocyte differentiation

SW22C6 is an induced pluripotent stem cell line bearing a p.G155fsX197 splice site mutation resulting in skipping of *CDKL5* exon 8 [37]. SW22 is the isogenic control line without the mutation. Both cell lines were a gift of Prof. David Millar at Cardiff University, UK. RET849 #13 iPSCs bearing a p.Glu55fs*74 frameshift mutation in *CDKL5* exon 5 and RET 849 #11, the isogenic control, were derived from patient fibroblasts by Prof. Ilaria Meloni and Dr. Alessandra Renieri, University of Siena, Italy [28]. RET849 lines were obtained from the lab of Prof. Simone Di Giovanni at Imperial College London, UK.

Non-tissue culture treated 6-well plates (Starlab, CC7672-7506) were coated with Vitronectin XF (StemCell Technologies, 07180) diluted to a final concentration of 10 µg/ml in CellAdhere Dilution Buffer (StemCell Technologies, 07183) and incubated for at least 1 hour prior to plating. iPSCs were thawed and maintained in mTeSR1 complete iPSC medium (StemCell Technologies), with a full medium change every day. When iPSC colonies covered 80% of well area and/or had dense centres in comparison to the edge of the colony, cells were dissociated by incubation with ReleSR (StemCell Technologies, 05872). Cells were then plated at a 1:10 split into Vitronectin XF coated six-well plates.

To differentiate the iPSCs into astrocytes, the cells were first used to generate Neural Progenitor Cells (NPCs) as an intermediate cell bank, via Neural Induction. Neural Induction was performed using SMAD inhibition Neural Induction kit (StemCell Technologies, #08581) according to manufacturer’s monolayer protocol. Cells were plated at a density of 2 - 2.5 x 10^5^ cells/cm^2^ in 6-well plates which had been earlier coated with 15 µg/ml Poly-L-Ornithine (Sigma-Aldrich, P4957) followed by 10 µg/ml Laminin (Sigma-Aldrich, L2020). Cells underwent a full medium change every day. After 6 to 9 days, cells were fully confluent and underwent passaging. Neural precursors were dissociated with Accutase (Capricorn Scientific, ACC-1B) and plated at a density of 1.5 - 2 x 10^5^ cells/cm^2^ in Neural Induction Medium supplemented with 10 µM Y-27632. Cells were passaged in the same manner once more before being plated into NPC Medium (StemCell Technologies, #05833). Neural Progenitor Cells (NPCs) were maintained in NPC Medium (StemCell Technologies, #05833). NPCs were plated at a density of 1.25 x 10^5^ cells per cm^2^ on 6-well plates coated with 15 µg/ml Poly-L-Ornithine (Sigma-Aldrich, P4957) and 10 µg/ml Laminin (Sigma-Aldrich, L2020).

Astrocytes were generated from NPCs described above according to the StemCell Technologies Astrocyte Differentiation Protocol and associated mediums (StemCell Technologies, #100-0013 and #100-0016). On astrocyte differentiation day 0, NPCs were plated at a density of 1.5 - 2x 10^5^ cells/cm^2^ into 6-well plates coated with 1% Geltrex Reduced Growth Factor Basement Membrane Mix (Fisher Scientific, 11612149) diluted in DMEM-F12 with 15 mM HEPES (Gibco, 11330032). 24 hours after passaging NPCs, NPC medium was replaced with Astrocyte Differentiation Medium (StemCell Technologies, #100-0013). Following astrocyte differentiation, immature astrocytes were plated into Astrocyte Maturation Medium (AMM; StemCell Technologies, #100-0016) supplemented with 10 µM Y-27632 (Merck, 688000). Immature astrocytes were plated onto Geltrex coated culture (as for astrocyte differentiation) at a density of 1.5 – 2 x 10^5^ cells/cm^2^. After 24 hours, medium was replaced with fresh AMM without Y-27632. After 19 - 23 in AMM, appropriate numbers of astrocytes were lysed for future analysis (protein, RNA, ICC – see sections below). A population of mature RET849#11 and RET849#13 astrocytes was maintained in culture for up to 31 subsequent days. During this time, cells were fed every 2 – 3 days and passaged when confluent.

### Mouse Astrocytes

All animal procedures were approved by the local ethical committee and performed in accordance with the United Kingdom Animals Scientific Procedures Act (1986) and associated guidelines. To study changes of hippocampal and cortical transcriptome C57BL/6J CRL inbred mice (Charles River) aged 3-4 months were used. Animals were group housed on a 12/12 dark/light cycle, and food and water were given *ad libitum*.

Five male Cdkl5^-/y^ mice and five male wild-type littermate controls [16] were sacrificed and brains dissected to remove the hippocampus and somatosensory cortex regions for use in astrocyte isolation and analysis. Harvested tissue was pooled according to genotype and stored in MACS Tissue Storage Buffer (Miltenyi Biotec, 130-100-008) overnight at 4 °C and subsequently dissociated the following day. Dissociation was performed according to Swartzlander et al. 2018 [38]. Miltenyi Myelin Removal Beads II (Miltenyi Biotec, 130-096-433) and LS columns (Miltenyi Biotec, 130-042-401) were subsequently utilised according to manufacturer’s instructions to remove myelin from cells. The flow through from myelin removal, containing unmyelinated cells suspended in MACS buffer, was collected and astrocytes were isolated using an Anti-ACSA-2 Microbead kit (Miltenyi, 130-097-679) and MS Columns (Miltenyi Biotec, 130-042-201) according to manufacturer’s instructions. The ACSA-2 positive cell population containing astrocytes and ACSA-2 negative cell population (consisting of other cell types) were subsequently counted using a haemocytometer, and RNA extracted using the Qiagen RNeasy RNA Extraction kit according to manufacturer’s instructions.

### AAV vectors

siRNA that efficiently knocked down CDKL5 was designed based on sequence scoring detailed by Fakhr et al. 2016 [39]. HEK293T cells were transfected in triplicate with 50 nM of either a candidate or control siRNA, using the Lipofectamine RNAiMAX Reagent (Invitrogen, 13778100) according to manufacturer’s instructions for a 24-well plate. 40 hrs post transfection, RNA and protein were extracted and analysed for CDKL5 expression.

A shRNA targeting CDKL5 was designed based on the most successful siRNA construct in knocking down CDKL5 in HEK293T cells (Supplemental Table 4). A loop sequence linking the sense and anti-sense siRNA strands was chosen based on successful shRNAs in the literature [40, 41]. A control construct with the Scrambled shRNA sequence was also designed. Following successful ligation into a gutless vector, CDKL5 shRNA and Scrambled shRNA sequences were under the control of a ubiquitous U6 promoter whilst eGFP (included as a marker for transduction) was under the CMV promoter. The resulting plasmids were used for AAV vector production with the AAV-DJ serotype plasmid and pXX6-80 adenovirus pHelper plasmid as standard protocols [28].

### Slice cultures

Human foetal cerebral cortical tissue aged between 15 to 18 post conception weeks (pcw) from terminated pregnancies was obtained with ethical consent from the MRC-Wellcome Trust Human Developmental Biology Resource (HDBR) under Project 200428. Ethical approval was obtained from the Newcastle and North Tyneside NHS Health Authority Joint Ethics Committee (REC reference: 23/NE/0135), and informed maternal consent was granted prior to collection of each sample. Human adult cortical tissue was obtained from anonymous patients undergoing neurosurgical tumour resection, approved by the County Durham & Tees Valley 1 Local Research Ethics Committee (06/Q1003/51) and with clinical governance approved by the Newcastle Upon Tyne Hospitals NHS Trust (CM/PB/3707), IRAS 173990. All patients gave informed consent for the use of such tissue for scientific research studies.

Adult and foetal human cortical brain tissue were collected and cultured according to methodology detailed in McLeod et al [42] with the following modifications for adult tissue. After sectioning, adult cortical slices were trimmed into small (0.5 cm x 0.5 cm) pieces and placed in a sterile submerged chamber containing oxygenated (95% O2/5% CO2) Hanks Balanced salt solution and 20 mM HEPES (305-315 mOsM and 7.3 pH) at room temperature (RT) for 15 minutes. Subsequently, slices were placed inside 6-transwell plates with 24 mm, 0.4μm pore polyester membrane inserts (3450; SLS) for 1 hour at 37°C, 5% CO₂ in ambient O₂ and 90% humidity. Each transwell contained 1ml of BrainPhys (5790; Stemcell Technologies) based culture media supplemented with 1 x N2, 1 x B27, 40 ng/ml Brain-derived Neurotrophic Factor, 20 ng/ml Glia-derived Neurotrophic Factor, 30ng/ml Wnt7a, 200 nM ascorbic acid, 1 mM dibutyryl cyclic AMP, 1 μg/ml laminin and 20 mM HEPES. Finally, the media was replaced for the final maintenance media without HEPES and changed every 2-3 days.

Slices were transduced by applying 3-5 µl of CDKL5 shRNA or Scrambled shRNA control AAV-DJ serotyped vector at 2.80 x 10^12^ vg/ml on to the top of each slice. All cultures were maintained for 2-3 weeks then subsequently fixed in 4% paraformaldehyde (PFA), flash frozen for RNA or protein extraction, or used for extracellular field potential recordings.

To investigate the extracellular excitability of foetal and adult human organotypic brain slice cultures, slices were placed in a recording chamber and continuously perfused at 34°C with oxygenated (95% O2, 5% CO2) normal (‘control’) artificial cerebrospinal fluid (ACSF) at a rate of 5-6 mls/minute. Multielectrode arrays (A16×1-2mm-100-177, NeuroNexus) were used to record extracellular field potentials connected to a RHD2000 (Intan Technologies) running Recording Controller software v2 (Intan Technologies) with output to a Micro1401 (Cambridge Electronic Design (CED)) and Spike2 software v8 (CED). A Hum Bug 50/60Hz noise eliminator (Digitimer) was also used to reduce background electrical noise (50 Hz) in Spike2 recordings.

Slices were left to equilibrate for 30 minutes before baseline recordings were acquired in normal ACSF for 10 minutes. Subsequently, slices were perfused with modified ACSF (normal ACSF replaced with 8 mM KCl and 0.25 mM MgSO4) to promote a state of hyperexcitability and further recordings were then acquired. Analysis of extracellular field potentials were performed offline in Spike2 using threshold detection protocols to quantify the frequency and amplitude of the events detected.

### Cellular Assays

#### Glutamate uptake assay

iPSC-derived astrocytes were plated at a density of 50,000 cells/well in a 96-well plate. WT iPSCs were additionally plated into a 96-well plate and allowed to grow until fully confluent. 24 hrs after plating, cells were washed twice with 200 µl warm HBSS, treated with 200 µM Glutamate (Sigma-Aldrich, G8415) and incubated at 37 °C for 4 hours. For glutamate release, fresh HBSS was added to the wells and cells were incubated for 1 hr at 37 °C. Subsequently, The Glutamate-Glo Assay Kit (Promega) was used to measure concentration of glutamate in cell culture supernatant according to manufacturer’s instructions. To calculate glutamate uptake, the amount of glutamate in the supernatant was subtracted from the amount originally added to the cells. Values for both uptake and release were normalised to protein concentration for the corresponding well. Protein was extracted from adherent cells using RIPA buffer to allow for normalisation to account for any variation in cell number in each well.

#### Cytokine Assay

To investigate response to stimuli of iPSC-derived astrocytes, cells were incubated with Tumor Necrosis Factor alpha (TNFα) and/or Interferon-gamma (IFNγ) and assessed for release of C-X-C motif chemokine ligand 10 (CXCL10) and Interleukin 6 (IL-6). To achieve this, astrocytes that had been in maturation medium for 49 days were plated at a density of 20,000 cells per well in a 96-well plate (Corning, 3595). The following day, varying concentrations of TNFα (Thermo Fisher Scientific, 300-01A) and/or IFNγ (Thermo Fisher Scientific, PHC4031) were added to the medium. After 24 hrs, culture supernatant was harvested and frozen at -20 °C. Cells were washed twice with cold phosphate-buffered saline (PBS) and underwent protein extraction with RIPA buffer, to check for any variation in cell number in each well. Protein concentration was quantified by BCA assay, as per manufacturer’s protocol. To quantify release of CXCL10 and IL-6, commercial Enzyme-Linked Immunosorbent Assay (ELISA) kits were utilised (Abcam, ab173194 and ab178013 respectively), according to manufacturer’s protocol.

#### Osmotic Assay

To investigate swelling dynamics of iPSC astrocytes, an osmotic challenge was performed. To achieve this, cells were plated in 8-well chamber slides (Thermo Fisher Scientific, 177402) at a density of 10,000 cells per well. The next day, medium was replaced with AMM medium diluted in water at 3 different concentrations: 100% medium, 90% medium, 75% medium and 50% medium. After 2 minutes, cells were washed once with PBS and fixed with PFA. Cells were stained with Rabbit anti-GFAP and mounted with ProLong Gold Anti-Fade Mounting medium with DAPI. After image acquisition using the Olympus BX63 Fluorescence Microscope, images were analysed using FIJI.

### Protein extraction and quantification

Protein extraction from cultured cells was performed directly in culture vessels, using RIPA buffer supplemented with 1:100 EDTA and 1:100 HALT Protease Inhibitor Cocktail according to manufacturer’s instructions.

Mouse brain tissue and human organotypic brain slices were snap-frozen and protein extracted using T-PER Protein Extraction reagent (Thermo Fisher Scientific) supplemented with 1:100 EDTA and 1:100 HALT Protease Inhibitor Cocktail according to manufacturer’s instructions.

Total protein concentration was quantified via the Bichronic Acid (BCA) Assay (Thermo Fisher Scientific, 23225) according to manufacturer’s instructions. The absorbance was read using a microplate reader at 540 nm and analysed using associated SoftMax Pro 5.4 software. Absorbance values for the standards were plotted against known concentrations to generate a standard curve. This was used to extrapolate protein concentration of each assayed sample.

### Western blot

Sodium dodecyl sulphate-polyacrylamide gel electrophoresis (SDS-PAGE) was performed using 4-15% Mini-PROTEAN TGX Precast protein gels (Bio-Rad, 4561084). 50 µl of protein sample was loaded into each well, or 20 µl Novex Sharp Pre-stained Protein Standard (Invitrogen, LC5800). Protein was transferred to a Immobilon-P transfer membrane (Millipore, IPVH00010) in the Mini Trans-Blot Electrophoretic Transfer Cell (Bio-Rad, 1703930). Membranes were blocked for 1-2 hrs at RT using 5% Milk in 0.1% PBST. Primary antibodies were diluted in blocking buffer and staining was conducted overnight with gentle rocking at 4 °C, according to concentrations in Supplemental Table 1. The next day, membranes were washed thrice in 5% milk for 20 minutes per wash, before being incubated in secondary antibody solution for 1 hr at RT (Supplemental Table 2). The membranes were then washed thrice in 0.1% PBST at RT. Finally, membranes were incubated in SuperSignal West Pico Chemiluminescent Substrate (Thermo Fisher Scientific, 34087) for at least 5 minutes, according to manufacturer’s instructions. Membranes were imaged using the GeneGnome XRQ Imaging System. Semi-quantitative protein expression analysis was performed by analysing the images using ImageJ, according to the protocol by Stael et al [43].

### DNA purification and analysis

DNA extraction from CDD patient iPSCs was performed using the Qiagen DNeasy Blood and Tissue DNA Extraction Kit according to the manufacturer’s protocol for cultured cells.

DNA fragments corresponding to the targeted region of the CDKL5 gene were amplified using PCR, with the Platinum Taq DNA Polymerase kit according to manufacturer’s instructions. PCR products were purified using the QIAquick Spin PCR Purification kit according to manufacturer’s instructions. Final elution of PCR products was performed using 50 µl nuclease-free water.

### RNA Extraction and RT-qPCR

RNA was extracted from cultured cells using the Qiagen RNeasy RNA Extraction Kits. For RNA extraction from < 5 x 10^5^ cells, or from human organotypic brain slice cultures, the RNeasy Micro Kit was used, whereas for RNA extraction from between 5 x 10^5^ and 1 x10^7^ cells the RNeasy Mini Kit was used. RNA quality was assessed using the Agilent Bioanalyser RNA Total Picochip assay and associated software. An RNA integrity number (RIN) was assigned to each sample based on parameters set out by the software default settings. RINs > 7.5 were acceptable for future sample analysis via qPCR and RNAseq.

Reverse transcription (RT) was performed using The Precision nanoscript2 RT kit was utilised according to manufacturer’s instructions (PrimerDesign/Novacyt) to generate cDNA for use in qPCR. Between 10 ng and 200 ng starting RNA was utilised depending on yield and downstream requirements. No reverse transcription (NRT) controls were generated with the same steps, except for the exchange of the reverse transcriptase enzyme with nuclease-free water.

Quantitative PCR (qPCR) was used to quantify the expression of several genes of interest. This was performed using the Brilliant II SYBRÒ Green qPCR kit according to manufacturer’s instructions. Wells were subsequently analysed using the Agilent Aria Mx qPCR machine and associated Agilent Aria 1.5 software. Gene expression analysis was performed via the Livak method (delta-delta Ct) [44].

### RNA Sequencing and Analysis

RNA samples were submitted to either the Imperial BRC Genomics Facility or Novogene, according to their respective sample submission guidelines, for subsequent library preparation and RNA sequencing. To summarise processes performed by the RNAseq facilities: mRNA isolation was first performed through polyA selection, then cDNA libraries were prepared via RNA fragmentation and reverse transcription, and resultant libraries quantified via Real-Time PCR. Sequencing was carried out using the NextSeq2000 with P3 flow cell and 200 cycles (Imperial BRC Genomics Facility) or the NovaSeq X Plus Series (PE150) (Novogene). Both sequencing methods produced paired-end reads with 150 bases per end. > 25 million reads were obtained for all samples. Quality control was performed either through FastQC [45] or fastp [46]. Fastq files were used as an input into Salmon and the reads were quasi-mapped to the relevant transcriptome according to the Salmon vignette [47]. The transcriptome sequence from GRCm39 was used for mouse samples, whereas GRCh38 was used for human samples (accessed from ftp.ensemble.org/pub, release 109). Salmon outputs were imported and formatted for downstream analysis using tximport [48].

Differential gene expression analysis for mouse samples was performed using NOIseq using the protocol in the vignette specifically aimed at samples without replicates (NOIseq-sim), using false discovery rate multiple correction testing [49]. Differentially expressed genes were extracted using a q value cut-off of 0.95 and lfc threshold of 1 and saved as .csv files. To analyse the GO terms that the differentially expressed genes mapped to, the ClusterProfiler R package was used according to the associated vignette [50], along with the Mus Musculus ensembl database (accessed via org.Mm.eg.db). Enriched GO terms were identified using the enrichGO function, with a p-value cut-off of 0.05 and a q-value cut-off also of 0.05. Moreover, the STRING database was used to investigate protein-protein interactions relating to differentially expressed genes [51].

All code for differential gene expression analysis of human samples was written based on the DESeq2 vignette [52]. Differential gene expression analysis was performed using the “DESeq” function with default parameters. Log fold change (lfc) shrinkage was performed using the ashr workflow [53]. Differentially expressed genes were extracted using an adjusted p-value (padj) value of < 0.05 and lfc threshold of > 1.5. Volcano plots were generated using the EnhancedVolcano package according to the relevant vignette [54]. Gene set enrichment analysis was performed as for mouse samples, using the enrichGO function of the clusterProfiler package [50]. Dot plots were generated using enrichplot [55]. Heatmaps were generated using pheatmap [56]. As with the mouse samples, the online STRING database was used to investigate protein-protein interaction networks associated with differentially expressed genes [51].

To investigate how the transcriptomic identity of cells in this study compared with those of other studies, datasets were compared with those in the literature. Publicly available datasets were accessed via the Gene Expression Omnibus (GEO) database and downloaded from the Sequence Read Archive as fastq files [57–72], for list of studies see Supplementary tables 3a and 3b. These files were then processed through the same Salmon-tximport-DESeq2 pipeline as described above. Single-cell and single-nuclei RNAseq datasets were analysed via publicly available count matrices, for a list of studies see supplemental table 3b [73–83].

Expression of genes of interest was recorded using tissue RNAseq data deposited at www.ebi.ac.uk/arrayexpress/experiments/E-MTAB-4840 [84] from 138 samples of human foetal cortical tissue taken at ages ranging from 7.5 to 17 PCW, and from various positions along the anterior posterior axis of the cortex including the temporal lobe.

### Immunostaining

To visualise protein expression in cultured cells, immunocytochemistry was performed. Cells were seeded in 8-well chamber slides at a density of 15,000 cells/well. 24 – 48 hrs after plating, cells were washed with DPBS (Sigma-Aldrich, D8537) before being fixed in 4% PFA. Cells were permeabilised by incubation with PBS/BSA/Triton Buffer for 10 minutes at RT. Wells were subsequently washed twice with 1x PBS and once with PBS/BSA (1%), before being blocked with PBS/BSA (1%) for 5 – 15 minutes at RT. Primary antibodies were diluted in 150 – 200 µl PBS/BSA (1%) per well according to concentrations in Supplementary table 1 and incubated in the dark at 4 °C overnight. The following day, wells were washed thrice with PBS/BSA (1%), before being incubated in the dark for 30 - 45 minutes with 150 – 200 µl secondary antibody solution prepared in PBS/BSA (1%) according to specified dilutions (Supplemental table 2). Finally, wells were washed 4 times with 1 x PBS and chambers removed according to manufacturer’s instructions. Glass coverslips were mounted on top using ProLong Gold Antifade Mounting Medium with DAPI (Invitrogen, P36931). Slides were imaged using an Olympus BX63 Fluorescence Microscope and images processed using ImageJ/FIJI. Images of stained cells were processed in QuPath, using the “Positive Cell Detection” tool. To quantify the number of branches per cell in mature iPSC-derived astrocytes, GFAP 20 x stained images were processed using skeletonization in FIJI.

Immunostaining of acute and organotypic human cortical slices was performed as described by McLeod et al. [42, 85]. Slides were imaged using a Zeiss LSM800 Airyscan Confocal Microscope and processed using ImageJ/FIJI.

### Statistical Methods

Statistical analysis was performed with GraphPad Prism 10.0. First, data normality was assessed using a Shapiro-Wilk test. For comparisons between the means of two groups, if data were normally distributed (Shapiro-Wilk p-value > 0.05) a Student’s t-test was performed; otherwise the non-parametric Mann-Whitney test was performed. Where the mean of three or more groups was compared, a one-way ANOVA was performed. When assessing the influence of more than one independent variable a two-way ANOVA was performed. Tukey’s post hoc multiple comparisons test was applied to ANOVAs in which a significant main effect or interaction was detected. Statistical significance was defined as *p* < 0.05 for individual comparisons, and adjusted *p* (padj) < 0.05 for multiple comparisons.

## Results

### Astrocytic expression of CDKL5 in human and mouse

The expression of CDKL5 in astrocytes at both the mRNA and protein level was first confirmed. Primary mouse and human astrocyte cell lines showed expression of CDKL5 protein (Figure 1a & 1b). Moreover, analysis of publicly available bulk RNAseq datasets revealed robust CDKL5 mRNA expression in isolated prenatal and postnatal cortical human astrocytes, and in astrocytes isolated from the hippocampus and cortex of juvenile and young adult mice (Figure 1c & 1d). Interestingly, expression of *CDKL5* was significantly lower in commercial primary astrocyte lines than astrocytes isolated from human brain tissue. Finally, analysis of publicly available single-cell RNAseq (scRNAseq) datasets revealed that a higher proportion of human astrocytes expressed *CDKL5* compared to mouse astrocytes (Figure 1e), indicating that the kinase may be more relevant to human astrocytes than mouse astrocytes.

**Figure 1.**
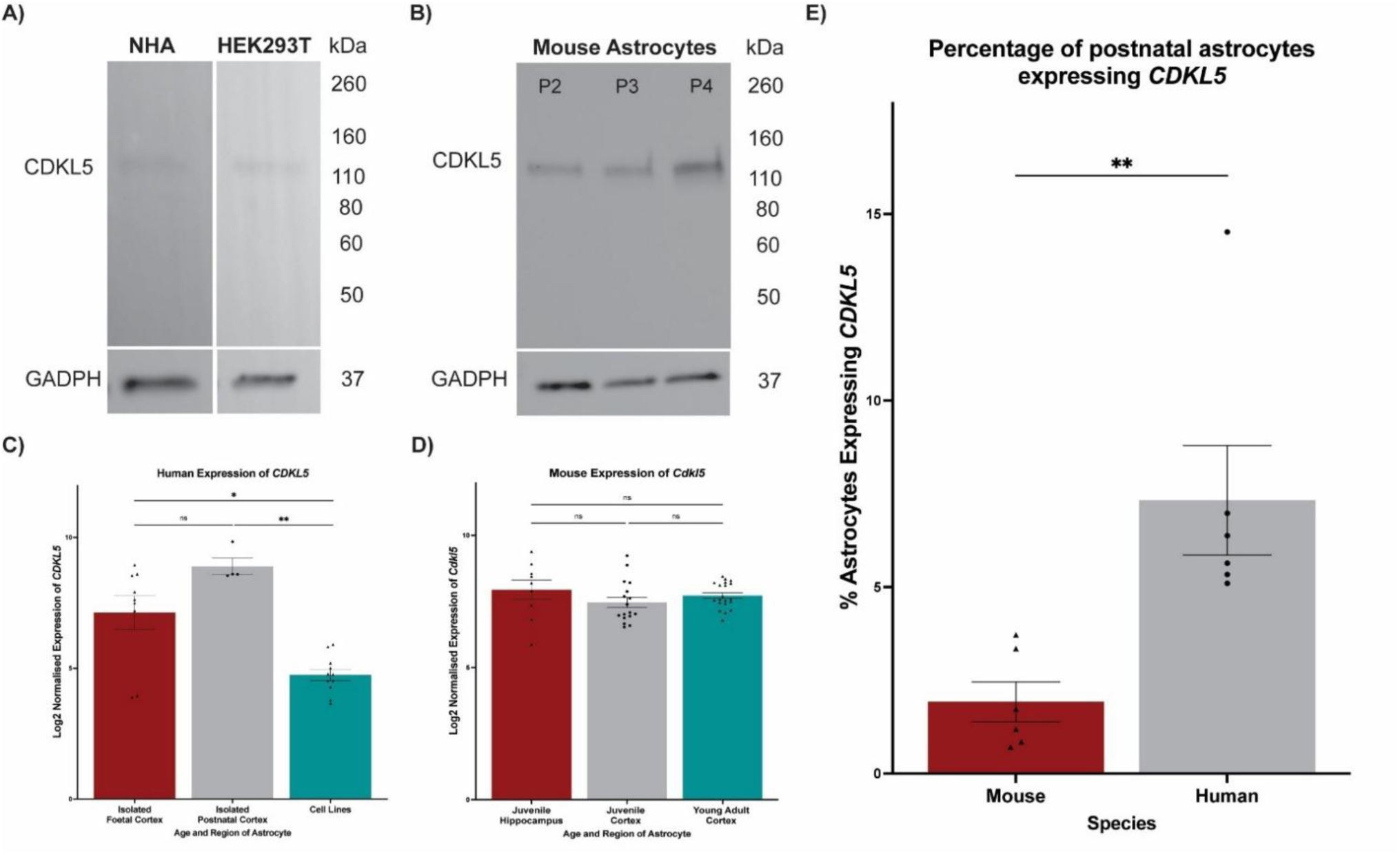
CDKL5 expression in mouse and human astrocytes. **A)** Western blot showing CDKL5 protein expression in Normal Human Astrocytes (NHAs), with HEK293T positive control. **B)** Western blot showing CDKL5 protein expression in primary mouse astrocytes (Sciencell) at three subsequent passages. **C)** Log_2_ normalised *CDKL5* transcript counts for human astrocytes in publicly available datasets. Isolated foetal cortex = astrocytes isolated from cortex of human foetal brains from elective pregnancy terminations, (n = 9 from 3 studies); Isolated postnatal cortex = as above but from human brain tissue at adolescent (10-18 yrs) or adult (19 yrs+) age (n = 4 from 1 study); Cell lines = commercial astrocyte cell lines (n = 11 from 3 studies); Significance determined via Mann Whitney test in GraphPad Prism, ns = p > 0.05, * = p < 0.05, ** = p < 0.005. Bars show mean, error bars are SEM. **D)** Log_2_ normalised *Cdkl5* transcript counts in mouse cortical and hippocampal isolated astrocytes from publicly available datasets. Juvenile Hippocampus = astrocytes isolated from hippocampi of mice aged P1-31 (n = 9 from 1 study); Juvenile Cortex = astrocytes isolated from the cortices of Juvenile (as defined above) mice (n = 17, total 5 studies); Young adult cortex = as described (n = 20, total 2 studies). Significance determined via one-way ANOVA with Tukey’s multiple comparisons, not significant (ns) = p > 0.05. Bars show mean, error bars are SEM. **E)** Percentage of astrocytes which had Cdkl5 expression in mouse and human astrocytes as determined in publicly available sc/snRNAseq datasets. n = 6 studies per species. Significance was determined using a Mann-Whitney test, p = 0.0022. Bars show mean, error bars are SEM.

### Differentiation of CDD patient iPSCs into astrocytes

Next, to interrogate the function of CDKL5 in human astrocytes, CDD patient iPSCs harbouring either a pGlu55*fs74 or pG155fsX197 C-terminal catalytic domain truncation of CDKL5 (loss of function) were differentiated into astrocytes, alongside isogenic controls obtained from the same individual patients (Figure 2a, Supplemental Figure 1). Following differentiation, iPSC-derived astrocytes showed robust expression of astrocytic markers, including GFAP, S100β, CD44 and ALDH1L1 (Figure 2 b-d). After 39 days in AMM, glutamate uptake was quantified for both the WT and CDKL5-mutant iAstros compared to an iPSC negative control. As expected, both the WT and CDKL5-mutant iAstros exhibited significantly increased glutamate uptake from the culture supernatant than the iPSCs, indicating that the glutamatergic homeostasis function of these cells was intact (Figure 2e).

**Figure 2.**
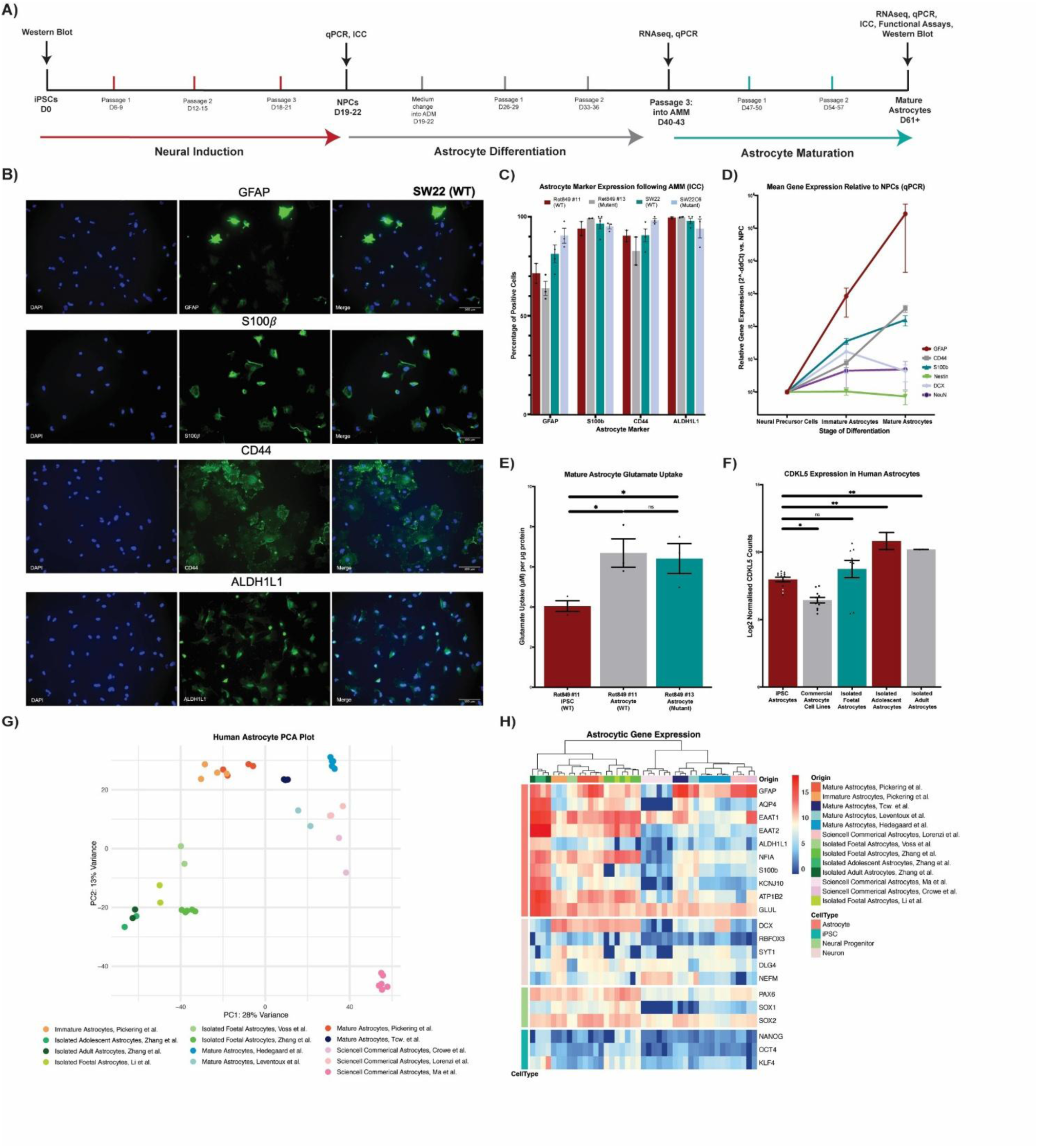
Differentiation of CDD patient-derived iPSCs into mature astrocytes. **A)** Schematic of CDD patient iPSC differentiation to astrocytes. **B)** Representative immunocytochemistry-immunofluorescence images of mature iPSC astrocytes. Staining shows GFAP, S100β, CD44, ALDH1L1 along with the nuclear stain DAPI. Scale bar represents 100 μm. **C)** Percentage of iPSC astrocytes expressing each marker, 10 x images (n=2 per cell line) were processed for Ret849 cell lines, whereas both 10 x (n=1 per cell line) and 20 x (n=2 per cell line) images were analysed for SW22 and SW22C6 cell lines. Error bars show SEM. **D)** RT-qPCR data showing relative gene expression throughout differentiation, compared to neural precursor cells (NPCs). **E)** Mean Glutamate Uptake in µM, per µg of total protein, for iPSCs and mature astrocytes (39 days in AMM, n = 3 separate wells per cell line). Cells were treated with 200 µM glutamate for 4 hours, glutamate uptake was calculated by subtracting the measured concentration of glutamate in the culture supernatant from concentration added. This was then normalised to total protein extracted from each well to account for differences in cell number. Two-tailed unpaired t-tests were performed to statistical significance. * = 0.01 < p < 0.05. Error bars are SEM. **F)** Log2 normalised counts of CDKL5 in iAstros (n=19), commercial human astrocyte cell lines (n=11) or isolated from human cortical tissue (foetal n= 9; adolescent n = 2; adult n = 2). Significance determined via a 2-way ANOVA with Tukey’s multiple comparisons. Ns = p > 0.05. * = p < 0.05. ** = p < 0.01. WT iAstros presented in this study are highlighted in white. **G)** PCA Plot comparing astrocyte RNAseq datasets publicly available in the literature (n=35) with those generated in this study. Each point represents a single sample, colour-coded depending on the study origin. Plot generated using DESeq2 and ggplot in R, using rlog transformation to stabilise variance across samples. **H)** Heatmap showing log_2_ normalised counts of genes associated with astrocytes, iPSCs, neural progenitor cells (NPCs) and neurons, in astrocytes generated through iPSC differentiation or isolated from human cortical tissue or commercial astrocytic cell lines. Origin refers to paper/source of each dataset, CellType refers to the cell type of which marker genes are associated. iAstros developed in the present research are referred to as Pickering et al. n = 43

Cell-type identity was further confirmed via transcriptomic comparison of immature and mature differentiated CDD astrocytes with that of acutely isolated, commercial primary and iPSC-derived astrocytes described elsewhere in the literature (Figure 2g & 2h). Transcriptomic analysis showed low expression of iPSC and neuronal genes in CDD patient iAstros, similar to differentiated astrocytes from other studies. A PCA plot showed that astrocytes from this study clustered together, and were on a similar PC2 axis as those from two other astrocyte differentiation studies. Astrocytes isolated from human tissue mostly clustered together and were closest on PC1 to the astrocytes in this study. Therefore, the CDD astrocytes in this study occupy an intermediate position between the isolated astrocytes and other differentiated astrocytes, and are more distinct from commercially available astrocytes.

*CDKL5* expression in iPSC astrocytes was confirmed by comparing the normalised counts of *CDKL5* across WT samples in this study and those in the studies listed above. As expected, *CDKL5* was expressed across all iAstros, including cells generated in the present study (Figure 2f). iAstros had significantly higher CDKL5 expression than commercial astrocyte cell lines, and significantly lower expression than isolated adolescent and adult astrocytes. There was no significant difference between CDKL5 expression in iPSC astrocytes and isolated foetal astrocytes (Figure 2f).

### Inflammatory dysregulation in CDD patient iAstros

To interrogate transcriptomic dysregulation between isogenic control (WT) and CDKL5-mutant iAstros, differential gene expression analysis was performed. A total of 50 differentially expressed genes were identified using DESeq2 with a threshold of padj < 0.05 and Log2 fold change (FC) > 1.5. Of these 50 genes, 16 were upregulated and the remaining 34 were downregulated (Figure 3a & 3b). Upregulated genes with the highest level of significance and Log2 FCs were the Major Histocompatibility Complex (MHC) Class II genes HLA-DRA (padj = 8.97E-16, log2 FC = 8.59) and HLA-DPA1 (padj = 1.22E-07, Log2 FC = 9.20), as well as the transcription factor NKX6-1 (padj = 1.37E-07, log2 FC = 4.43), which is normally expressed at low levels in astrocytic nuclei. Downregulated genes with the most significant padj and largest Log2 FC included the protocadherin PCDHGA7 (padj = 1.82E-24, Log2 FC = -6.71), ZNF736 (padj = 1.06E-4, log2 FC = -7.77) and the crystallin CRYBB2 (padj 1.06E-4, log2 FC = -5.73). Notably, the epilepsy-linked water channel AQP4 was significantly upregulated in CDKL5-mutant iAstros. Functional pathway enrichment revealed association of differentially expressed genes with MHC Class II protein complex and activity. Morover, STRING protein-protein interaction network analysis of differentially expressed genes identified 3 significant local network clusters, all of which were concerned with MHC Class II.

**Figure 3.**
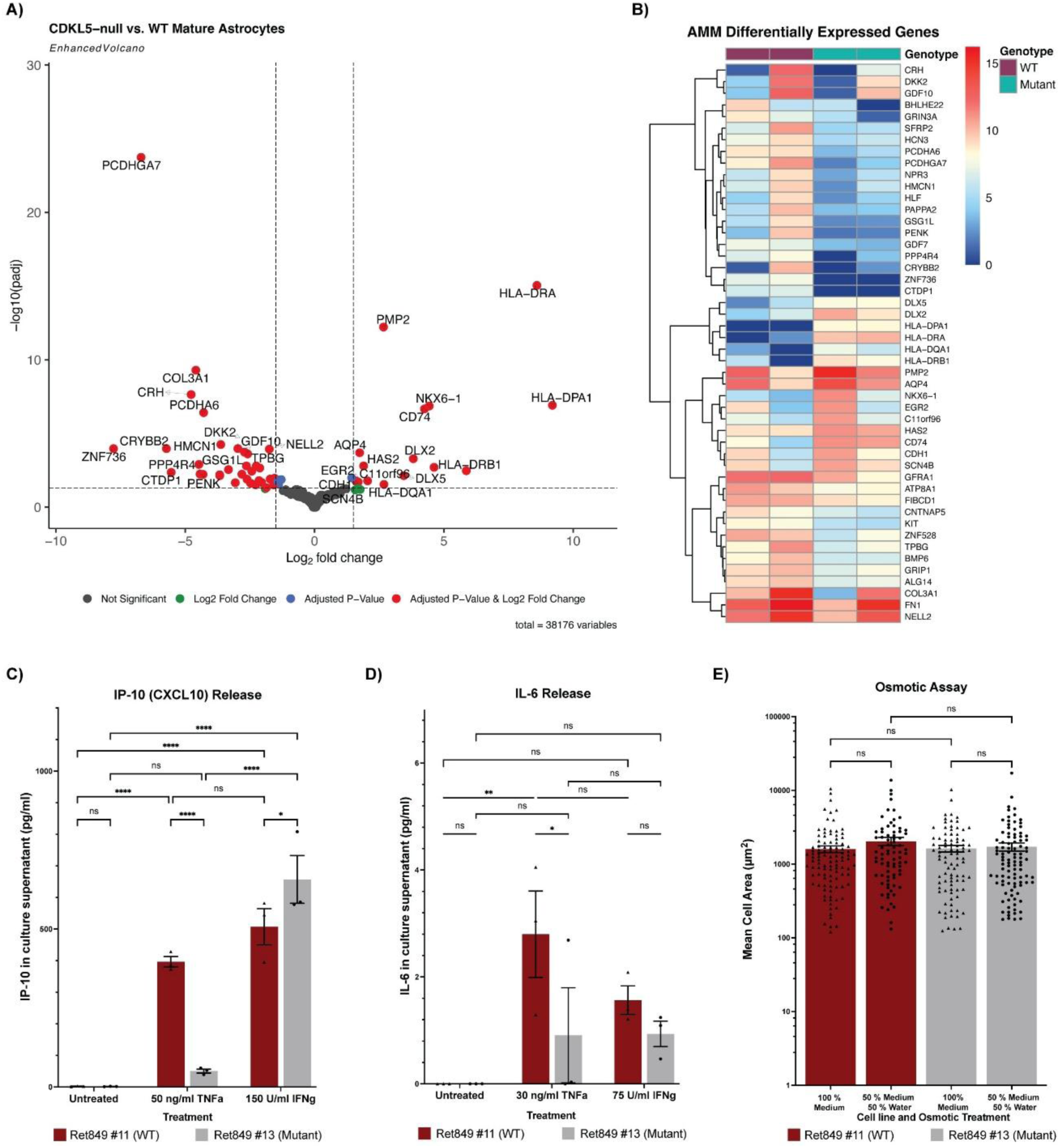
CDKL5-null astrocytes have transcriptomic and functional inflammatory deficits. **A)** Volcano plot showing -log10(adjusted p-value) against log_2_ fold change (lfc) of genes expressed by mature astrocytes generated in this study. Each dot represents a gene, red dots represent significantly differentially expressed genes between CDKL5-null (mutant) and WT astrocytes. Lfc threshold >= 1.5, padj < 0.05. n=2 samples per genotype. **B)** Heatmap showing log2 +1 transformed normalised counts of DEGs. **C)** Mean concentration of IP-10 (CXCL10) in WT and CDKL5 mutant iAstro culture supernatant (pg/ml), the day after: no treatment; addition of 50 ng/ml TNFa to culture medium; addition of 150 U/ml IFNg to cell culture medium. **D)** Mean concentration of IL-6 in WT and CDKL5-null (Mutant) iAstro culture supernatant (pg/ml), the day after: no treatment; addition of 30 ng/ml TNFα to culture medium; addition of 75 U/ml IFNγ to cell culture medium. For both C) and D), n = 3 culture wells per line. Statistical significance was determined using a two-way ANOVA with Tukey’s multiple comparisons. Ns = not significant. * = padj < 0.05, ** = padj < 0.01, *** = padj < 0.005, **** = padj < 0.001. Error bars show SEM. **E)** Mean cell area, in µm^2^, of WT and CDKL5-null iAstros exposed to normal AMM conditions (WT n = 104 cells, CDKL5-null = 91 cells) compared to those exposed to AMM diluted 50% in water for 2 minutes (WT n = 78 cells, CDKL5-null = 99 cells). Significance was tested using a two-way ANOVA with Tukey’s multiple comparisons. Ns = not significant. Error bars show SEM.

Given the immune-related dysregulation indicated in the transcriptomic analysis, cytokine and chemokine release in response to stimuli was investigated. To this end, iAstros were treated with either TNFα or IFNγ, and then the next day the concentration of CXCL10 and IL-6 in the culture supernatant was measured by ELISA. No significant differences were seen between total intracellular protein of WT vs. CDKL5-mutant iAstros within each treatment. As would be expected from functional astrocytes, both WT and CDKL5-mutant iAstros released IL-6 and CXCL10 in response to TNFα or IFNγ treatment (Figure 3c & 3d). However, the level of IL-6 released was very low, and only significantly more than the untreated iAstros when WT iAstros were treated with TNFα. CDKL5-mutant iAstros did not release significantly more IL-6 than their untreated counterparts under either treatment condition. Interestingly, the CDKL5-mutant iAstros had a significantly decreased release of both CXCL10 and IL-6 compared to WT iAstros in response to treatment with TNFα. Conversely, CXCL10 release was significantly increased in CDKL5-mutant iAstros compared to WT following IFNγ treatment (Figure 3c). Therefore, there appears to be a divergent response to stimuli in CDKL5-mutant iAstros compared to WT iAstros, particularly in terms of TNFα release.

To deconstruct the consequences of AQP4 dysregulation in iAstros, an osmotic assay was performed in which AMM culture media was diluted with water (50 % AMM and 50 % water) and cells were exposed to diluted medium for 2 minutes. iAstros were then immediately fixed and stained with GFAP to investigate the effect of the disrupted osmotic balance on cellular swelling. In both WT and CDKL5-mutant iAstros there was an increase in cellular area in response to dilution of AMM with 50% water, but neither increase reached the threshold of statistical significance (Figure 3e). Nonetheless, the WT cells appeared to have a slightly larger increase in area than the CDKL5-mutant cells following hypotonic stress, without reaching statistical significance. No difference in area was detected between the two genotypes in normal AMM conditions (100% AMM).

### Cytoskeletal dysregulation in CDD patient iAstros

Given that cytoskeletal organisation is a key role of CDKL5 in neurons, it was important to establish whether this function is conserved in astrocytes. As such, phosphorylation of a key neuronal cytoskeletal target of CDKL5, End-Binding protein 2 (EB2), was quantified. Interestingly, analysis of the Ret849 #11 and #13 mature astrocytes at three subsequent passages revealed that whilst the level of phosphorylation of EB2 remained constant, more EB2 was present in CDKL5-mutant cell lines, resulting in a significant decrease in the ratio of pEB2/EB2 (Figure 4a-d). The morphological properties of iPSC astrocytes were also determined via staining with GFAP and analysis with FIJI. As described above, there was no significant difference in cell area between WT and CDKL5-mutant iAstros. However, branching analysis showed that CDKL5-mutant iAstros had significantly more branching than WT iAstros, with an average of 7.1 branches per astrocyte compared to 5.4 in WT cells (Figure 4e-g).

**Figure 4.**
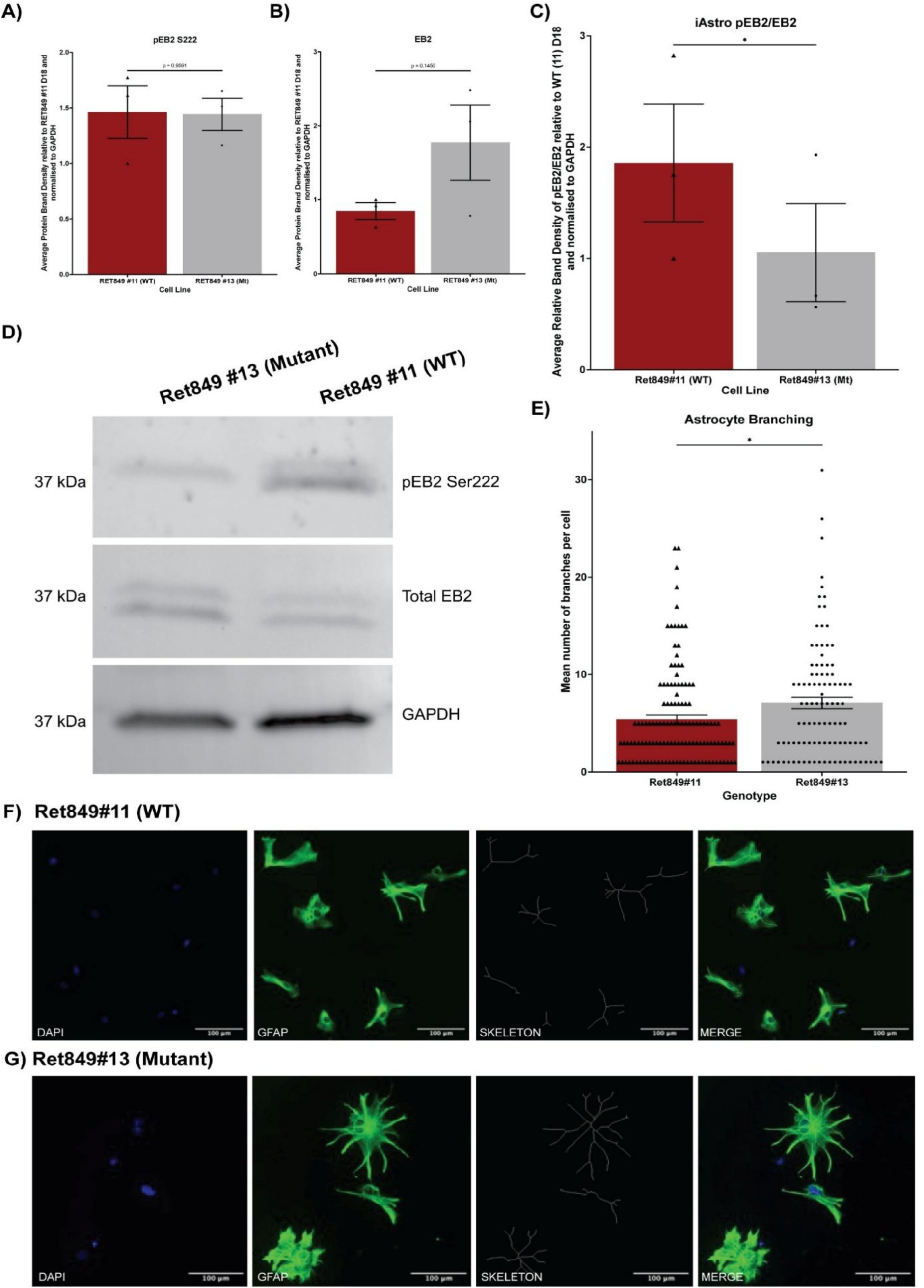
Cytoskeletal deficits in CDKL5-null iPSC astrocytes. **A)** Semi-quantitative analysis of protein band density of phosphorylated EB2, normalised to GAPDH and WT D18 sample. Analysis performed using n=3 separate time points during AMM to generate replicates from Ret849#11 and Ret#84913 cell lines. Time points were Day 18 (completion of maturation), Day 46 and Day 51. Time points were arbitrarily selected based on available cell-numbers for protein extraction. Significance assessed with two-tailed paired t-test. **B)** As for B but EB2. **C)** Ratio of pEB2 to EB2 following GAPDH normalisation in both Ret849#11 and Ret849#13 cell lines, using data represented in B) and C). A one-tailed paired t-test was used to determine whether there was a significant decrease in pEB2/EB2 ratio between WT and CDKL5-null (Mt) cells at each time-point. * represents statistical significance, p = 0.0424. Error bars are SEM. **D)** Representative western blot images of staining conducted on CDKL5-null (Ret849#13) and WT (Ret849#11) mature astrocytes following AMM protocol. Representative blot is from day 51. **E)** Mean number of branches per cell for WT and CDKL5-null iPSC astrocytes. Significance was determined using Mann-Whitney test, p = 0.041. Error bars show SEM. WT n = 127 cells, CDKL5-null n = 102 cells. **F)** Representative images of GFAP staining and subsequent skeletonization for branch counts in FIJI. Scale bar represents 100 μm in WT iAstros. **G)** As for F) but CDKL5-null iAstros.

Taken together, the use of iPSC-derived astrocytes as a model of CDD highlighted transcriptomic and functional deficits in CDKL5-mutant iAstros which warranted further investigation in a multicellular model of the disorder.

### Transcriptomic investigation of cortical and hippocampal mouse astrocytes

Next, we therefore sought to investigate CDKL5-null astrocytes in a mouse model of CDD. However, mouse astrocytes isolated from cortex and hippocampus had lower CDKL5 expression than human derived astrocytes (as in Figure 1). Subsequent interrogation by RNAseq did reveal significant transcriptomic differences such as cytoskeletal dysregulation between WT and CDKL5 KO astrocytes but no widespread changes in inflammatory response were observed (Supplemental figure 2). Moreover, whilst transcriptomic analysis showed a high level of expression of astrocyte markers suggesting a predominantly astrocytic cell population, many of the differentially expressed genes were not astrocyte-specific, suggesting that Cdkl5 deficiency does not lead to unique dysregulation in these cells in mice (Supplemental Figure 2). The lack of spontaneous seizures in this mouse model characterised in the literature [16], coupled with the lower level of astrocytic CDKL5 expression and relevance apparent in the present data, suggests that mice are a less useful model of astrocytes in CDD.

### Novel *in vitro* model of CDD using human organotypic brain slice cultures

Whilst interrogation of patient-derived iAstros offers insight into inflammatory and cytoskeletal deficits in CDKL5-mutant astrocytes, investigating network connectivity is required to better understand the full range of astrocytic functions of CDKL5. Therefore, a new model of CDD was generated using shRNA-CDKL5 AAV knockdown of CDKL5 in human foetal and adult organotypic brain slice cultures. First, the basal expression of *CDKL5* in foetal tissue was confirmed using an RNAseq data from Lindsay et al. [84], which showed a gradual increase during prenatal development up until pcw 12, at which point expression began to plateau (Figure 5a). This suggests that CDKL5 may be functionally relevant in the cortex during development.

**Figure 5.**
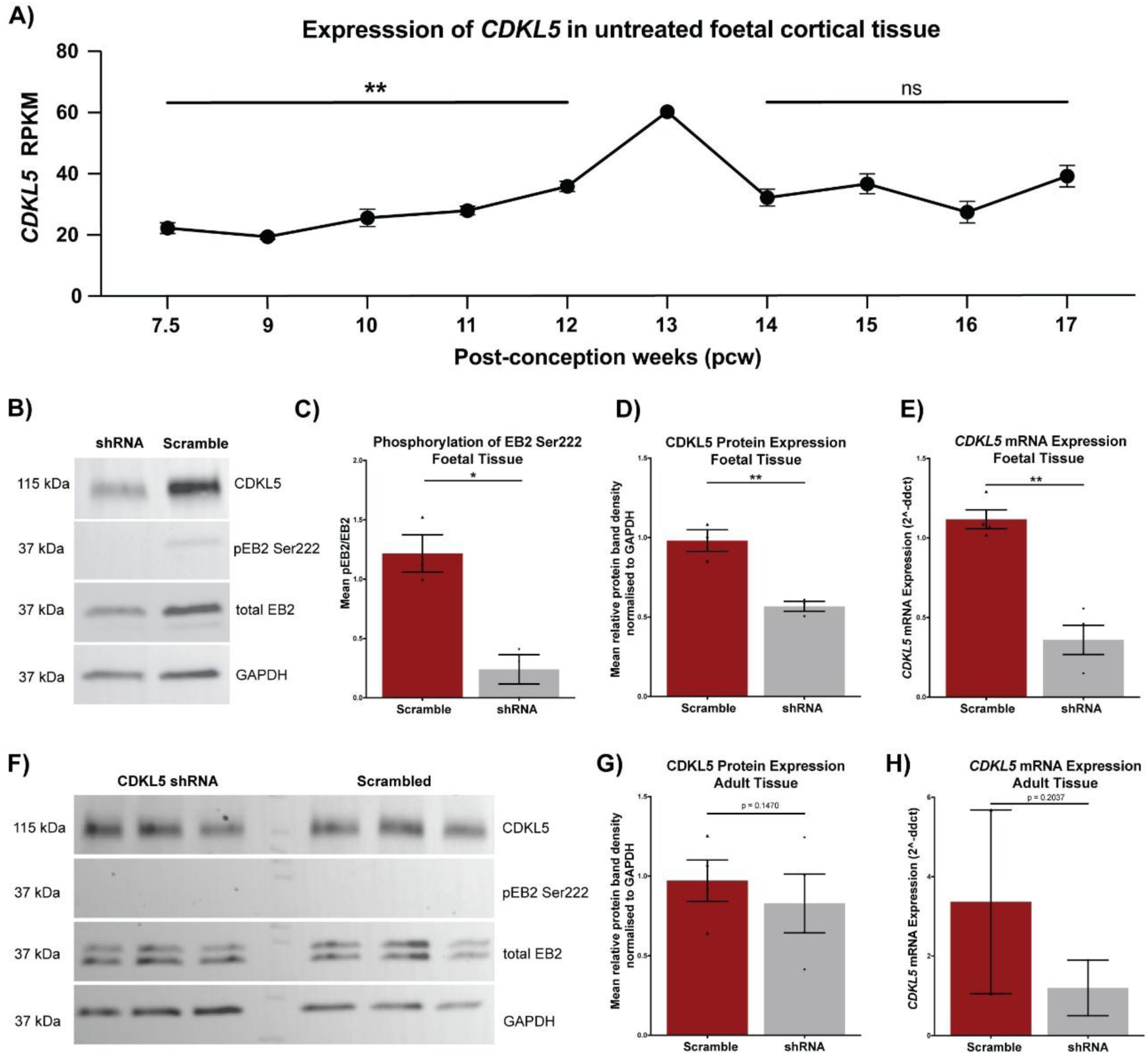
Proteomic analysis of CDKL5 knockdown in foetal and adult cortical slices. **A)** Reads per Kilobase per Million mapped reads (RPKM) of CDKL5 in WT foetal cortical tissue at various post-conception weeks (pcw). Statistical significance determined using one-way ANOVA with Tukey’s multiple comparisons. ** = padj < 0.005. Error bars are SEM. **B)** Representative western blot staining of transduced foetal tissue protein, for CDKL5, pEB2, EB2 and GAPDH. **C)** Ratio of pEB2/EB2 in transduced foetal cortical tissue, following normalisation to GAPDH. Significance determined by paired t-test. **D)** Semi-quantitative analysis of CDKL5 protein band density for transduced foetal cortical tissue treated with Scrambled shRNA or CDKL5 shRNA. Significance determined via paired one-tailed t-test. n=3 biological cases with 3 technical replicates per case (n=9 slices per condition). p=0.0056. Error bars are SEM. **E)** RT-qPCR analysis of CDKL5 mRNA expression in cortical foetal slices treated with either CDKL5 shRNA or Scrambled shRNA. n=4 biological cases with 2-3 technical replicates per case (n=11 slices per condition). Significance determined via paired one-tailed t-test. p=0.0014. Error bars represent SEM. **F)** Representative western blot staining of transduced adult tissue protein, for CDKL5, pEB2, EB2 and GAPDH. Protein samples from three separate slices per vector, originating from a single biological case, are shown. **G)** As for D but transduced adult cortical tissue, n=4 biological cases with 2-3 technical replicates/case (11 slices/condition). **H)** As for E but transduced adult cortical tissue, n=2 biological cases with 2-3 technical replicates/case (5 slices/condition).

A novel shRNA targeting CDKL5 delivered via an DJ serotyped AAV vector exhibited robust transduction in these cultures, as determined by the co-expression of eGFP in both neurons and astrocytes (Supplemental Figure 3 and 4). In foetal tissue, this elicited a 42% knockdown of CDKL5 protein compared to Scrambled shRNA treated slices (Figure 5d). This was combined with an 80% reduction of pEB2/EB2 in shRNA-CDKL5 AAV treated foetal slices (Figure 5c). In adult cortical slices the degree of CDKL5 protein knockdown was lower, at just 16% (Figure 5g). Moreover, no phosphorylated EB2 at Ser222 was detected in the adult cortical organotypic brain slice cultures, despite total EB2 detection (Figure 5f).

Finally, to confirm functional effects of CDKL5 knockdown and show proof of concept that this model has a hyperexcitable phenotype, the extracellular network activity was assessed. Two different solutions were used to bath the slices in during recording – normal artificial cerebral spinal fluid (ACSF) followed by modified ACSF. Normal ACSF is designed to recapitulate basal conditions and enable baseline recordings, whereas modified ACSF has high potassium and low magnesium, to promote a state of hyperexcitability in the slices. For both foetal and adult organotypic cultures, in slices transduced with Scrambled shRNA AAV there was a trending increase in the frequency and amplitude of events (i.e. transient increases in extracellular field potentials), between the time-period spent in normal (control) ACSF, and the time-period spent in modified ACSF (Figure 6). This is indicative that the modified ACSF is functional in terms of promoting a state of hyperexcitability in the slices. However, the slices transduced with the CDKL5 shRNA showed a similar frequency of events in the normal ACSF as the Scrambled shRNA did in the modified ACSF, implying that there was an increase in events in normal ACSF upon CDKL5 knockdown. There was also a trending increase in the same manner in terms of amplitude of events. There was no notable difference in frequency of events between normal and modified ACSF in slices treated with CDKL5 shRNA, in contrast to the recordings of the Scrambled shRNA treated slices. This could be indicative of the CDKL5 shRNA slices reaching peak hyperexcitability in normal ACSF. Therefore, this data suggests that there is an increase in spontaneous network hyperexcitability in CDKL5 shRNA vector treated slices, arising from knockdown of CDKL5 in both neurons and astrocytes. This finding was consistent across foetal and adult organotypic brain slice cultures (Figure 6).

**Figure 6.**
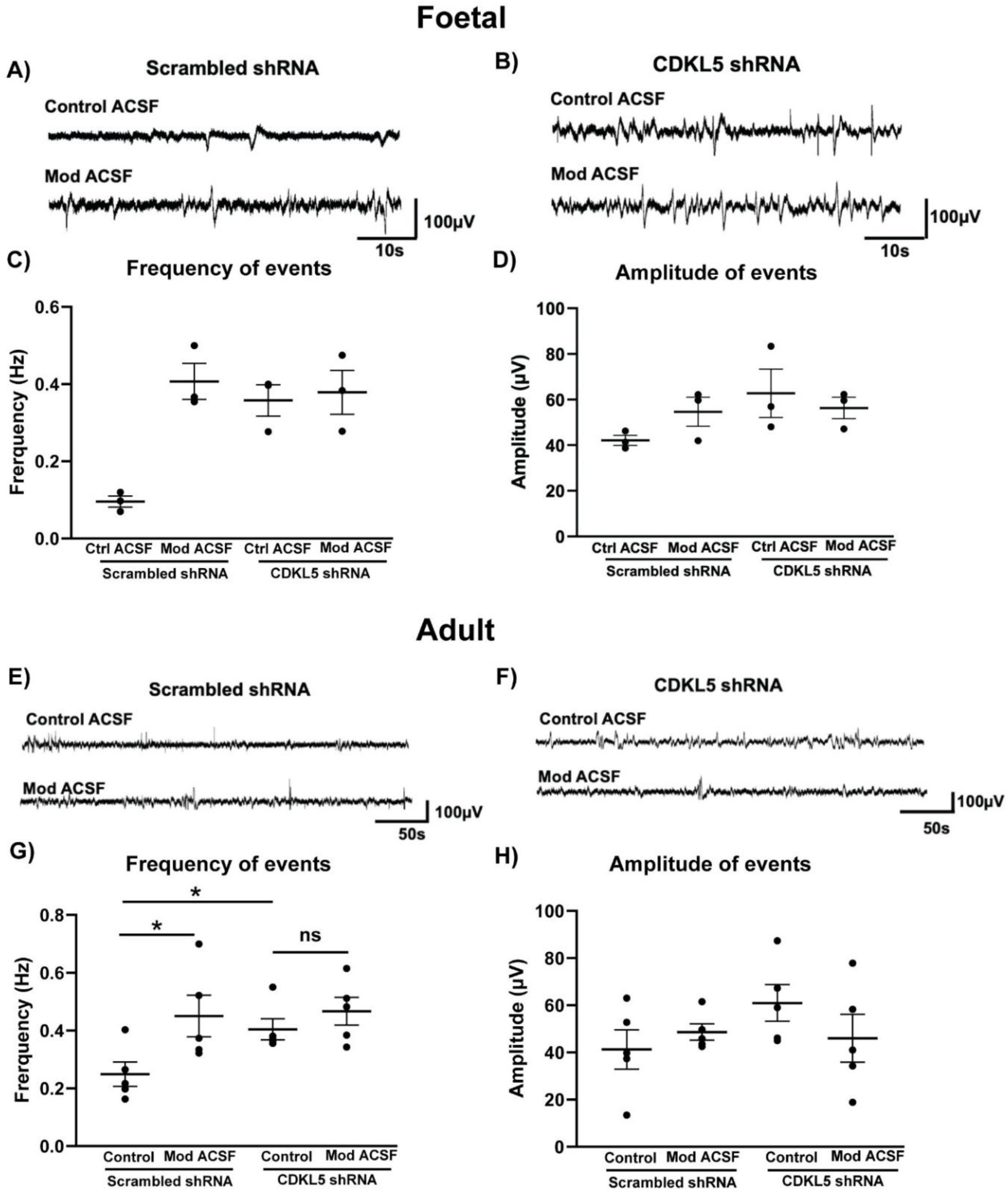
Functional analysis of CDKL5 knockdown in foetal and adult cortical slices. **A)** Representative traces of extracellular recordings for foetal tissue transduced with Scrambled shRNA. ACSF = artificial cerebral spinal fluid. Mod ACSF= modified ACSF, with high K^+^ and low Mg^2+^, to promote a state of hyperexcitability. **B)** Representative traces of extracellular recordings for foetal tissue transduced with CDKL5 shRNA. **C)** Frequency of spike events in control (ctrl) ACSF and Mod. ACSF for transduced foetal tissue. n=1 case with 3 slices/condition. No statistics were performed due to lack of biological replicates. Error bars represent SEM. **D)** Amplitude of spike events in control ACSF and Mod. ACSF for transduced foetal tissue. n=1 case with 3 slices/condition. Error bars represent SEM. **E)** Representative traces of extracellular recordings for adult tissue transduced with Scrambled shRNA. **F)** Representative traces of extracellular recordings for adult tissue transduced with CDKL5 shRNA. **G)** As for C) but for transduced adult tissue. n=2 cases with 5 slices/condition and significance determined via a two-way ANOVA (p < 0.05). Error bars represent SEM. **H)** As for D) but for transduced adult tissue. n=2 cases with 5 slices/condition. No statistically significant differences were found between conditions.

### Investigation of astrocyte function in CDKL5 shRNA treated human organotypic brain slice cultures

AQP4 expression was investigated via immunohistochemistry staining of 17 pcw foetal occipital cortex (Figure 7a), in which large AQP4+ GFAP+ cells with multiple emanating branches were identified. An RNAseq dataset corresponding to foetal tissue from 7.5 pcw all the way through to 17 pcw was next analysed for *GFAP* and *AQP4* expression ([84], Figure 7b and c). *GFAP* expression remained relatively consistent throughout the prenatal timepoints, indicative of the high expression in radial glial cells during this period. *AQP4* expression was considerably lower than *GFAP*, remaining below 1 RPKM until a rapid increase between 11 pcw and 12 pcw and reaching a peak of 23.03 RPKM at 16 pcw.

**Figure 7.**
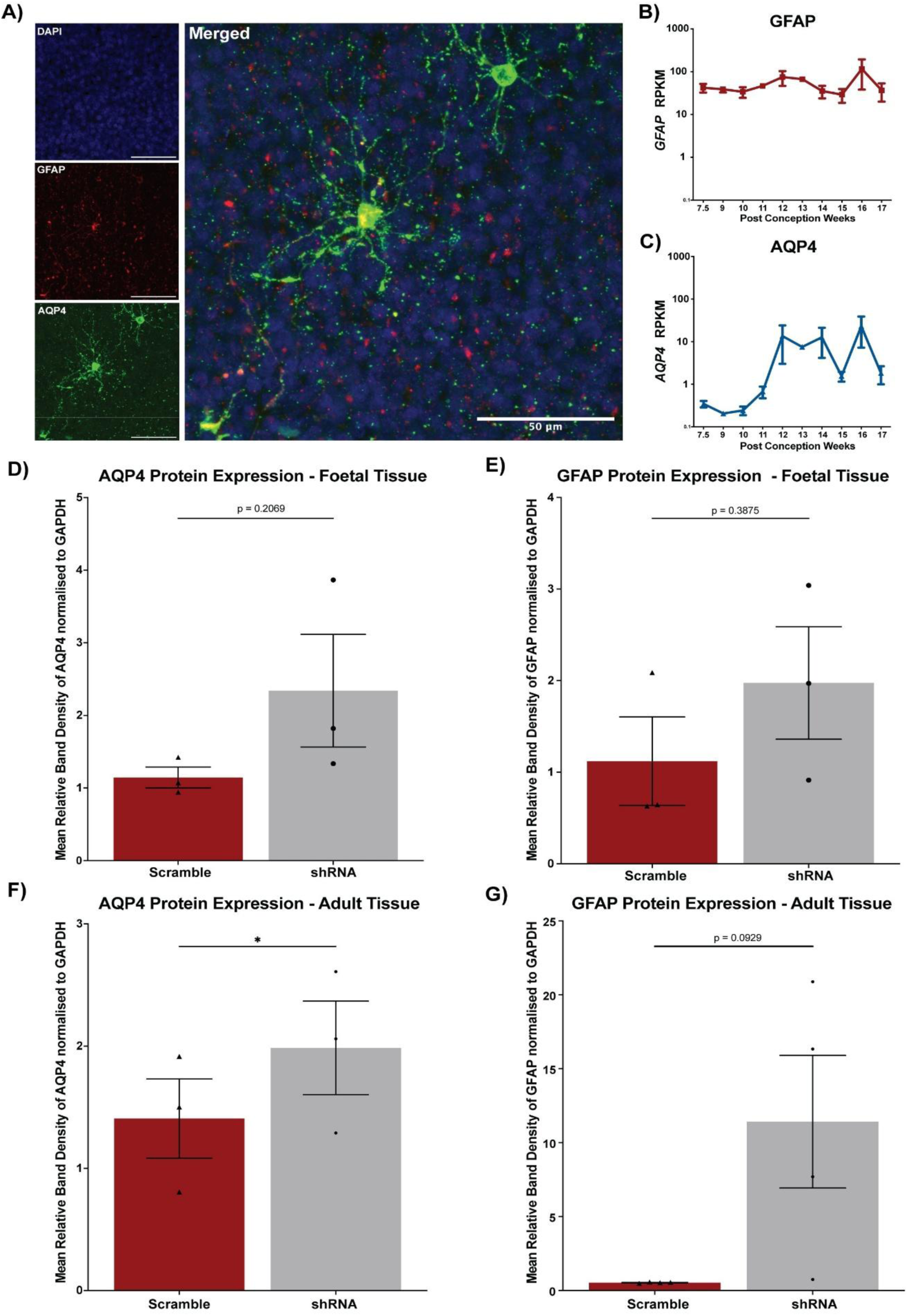
Astrocyte-specific AQP4 is expressed in foetal cortical tissue and is dysregulated following CDKL5 knockdown. **A)** Representative immunofluorescence staining for AQP4 (green), GFAP (red) and DAPI (blue) in WT foetal pcw 17 occipital cortex. Scale bar represents 50 µm. **B)** Reads per Kilobase per Million (RPKM) of *GFAP* in foetal cortical tissue at various post-conception weeks (pcw). No significant differences between ages were found (one-way ANOVA with Tukey’s multiple comparisons). Error bars are SEM. **C)** Reads per Kilobase per Million of *AQP4* in foetal cortical tissue at various pcw. No significant differences between ages were found (one-way ANOVA with Tukey’s multiple comparisons). Error bars are SEM. **D)** Semi-quantitative analysis of AQP4 protein band density for transduced foetal cortical tissue (Scrambled or CDKL5 shRNA). Significance determined via paired two-tailed t-test. Error bars are SEM. n=3 biological cases with 3 technical replicates per case (n=9 slices per condition). **E)** Semi-quantitative analysis of GFAP protein band density for transduced foetal cortical tissue (Scrambled or CDKL5 shRNA). Significance determined via paired two-tailed t-test. Error bars are SEM. n=3 biological cases with 3 technical replicates per case (n=9 slices per condition). **F)** As for D but adult tissue. n=4 biological cases with 2-3 technical replicates/case (11 slices/condition). **G)** As for E but adult tissue. n=4 biological cases with 2-3 technical replicates/case (11 slices/condition).

Given that AQP4 was upregulated in CDKL5-null iAstro transcriptomic datasets, the expression of this epilepsy-linked water channel was investigated in the foetal and adult organotypic brain slice cultures. Both AQP4 and GFAP expression were increased, albeit to varying extents, in CDKL5 shRNA treated foetal and adult slices compared to Scrambled shRNA (Figure 7d-g). It thus appears that an observed trend of increased AQP4 is consistent across all human models of CDKL5 deficient astrocytes presented in this study, but doesn’t reach significance in all models.

## Discussion

Given the role of astrocytes in synaptic development and homeostasis, we sought to a) develop and evaluate the utility of various models for the study of CDD astrocytes, and b) identify potential areas of astrocytic deficits in CDD. Initially, astrocytic CDKL5 expression was investigated, finding that whilst both mouse and human astrocytes express low levels of CDKL5, expression was higher in human astrocytes. We found that astrocytic CDKL5 appeared to be involved in the regulation of signalling pathways related to reactivity, cellular adhesion and cytoskeletal organisation in CDD patient iPSC-derived astrocytes. Changes to regulation of these pathways led to altered morphology and response to immunological stimuli. After subsequently determining that mice have limited value for study in this context due to lack of phenotypic recapitulation and very low astrocyte expression of CDKL5, we moved to develop a multicellular *in vitro* human model of CDD. This model allowed us to identify astrocyte-specific dysregulation (such as the upregulation of AQP4) and investigate the impact of a global CDKL5 knockdown on network activity. As such this study provides first proof-of-concept for a valuable new model with network connectivity that can be utilised to study CDD pathogenesis and develop therapeutic interventions.

### Cdkl5 in subpopulations of astrocytes in the mouse

We found no evidence that mouse astrocytes are devoid of CDKL5 expression. Data published by Batiuk et al. shows that only a very small proportion of mouse astrocytes express CDKL5, which could explain why some studies failed to detect any expression [76]. However, Batiuk et al. also show varied levels of expression of CDKL5 across different astrocytic subtypes. Not only is this another explanation for some studies finding no mouse astrocyte CDKL5 expression, but it also offers insight into what functional relevance CDKL5 may have in these cells. The subtype with the highest proportion of CDKL5 -expressing astrocytes was AST3, closely followed by AST2 and AST4 [76]. Batiuk reported AST2 and AST3 astrocytes as having little or no GFAP expression, whilst AST1 – of which just 0.30% of cells were Cdkl5-positive – had the highest expression of GFAP [76]. Therefore, GFAP and Cdkl5 are not substantially co-expressed in all mouse astrocytes and the use of GFAP-Cre mouse knockouts may not accurately report on the astrocytic function of *Cdkl5* [86]. Interestingly, interrogation of scRNAseq datasets showed that a higher proportion of human astrocytes expressed *CDKL5* than mouse astrocytes. Furthermore, whilst cytoskeletal regulation appeared to be impacted by Cdkl5 deficiency in isolated mouse astrocytes, the very low mapping of differentially expressed genes to astrocyte-specific processes calls into question the contribution of these cells to the mouse CDD phenotype. Therefore, it is possible that astrocyte function is more affected by CDKL5 deficiency in humans than in current mouse models of the disorder.

### Transcriptomic and functional deficits in iAstros generated from CDD iPSCs

We explored basal CDKL5 expression in human astrocytes confirming previous observations in primary human astrocytes noted by Gao et al. [28]. We then demonstrated the successful differentiation of patient iPSCs into astrocytes, representing proof-of-concept for the generation of astrocytes from CDD patient cell lines. Importantly, these iPSC astrocytes express CDKL5 at a higher level than commercial astrocytes lines and at a similar level to isolated human foetal astrocytes, making them an ideal model for the study of astrocytic dysfunction in CDD.

We investigated the differences between CDD iAstros and isogenic controls, finding multiple avenues of dysregulation. Several MHC Class II related genes were upregulated in the CDKL5-mutant iAstros, suggesting an immunological element to astrocytic dysfunction in CDD. The upregulation of MHC II genes can be attributed in astrocytes to the interferon-gamma inducible transactivator CIITA [87]. Whilst the expression of CIITA (encoded by the MHC2TA gene) was not significantly altered in the present dataset, it has been well reported in the literature that CIITA activity is downregulated by its phosphorylation [88, 89]. Specifically, CIITA phosphorylation is thought to inhibit the transcriptional activation of MHC II genes [90]. One motif at which such phosphorylation occurs is Ser286, which matches that of the CDKL5 phosphorylation consensus motif (RPGS, [19]), suggesting that CDKL5 could be capable of phosphorylating CIITA at this site. In this case, a loss of functional CDKL5 would therefore result in increased MHC II transcription. Another mechanism through which CDKL5 could affect the activity of CIITA is through ERK1/2, which is a known CDKL5 target and has also been shown to directly phosphorylate CIITA at Ser286, Ser288 and Ser293 to inhibit MHC II upregulation [90, 91]. Therefore, it is possible that CDKL5-null astrocytes have reduced phosphorylation of CIITA through one of these two mechanisms, causing upregulation of MHC II genes and altered reactivity. However, further studies would be required to determine whether this occurs in CDKL5-deficient astrocytes.

Having established the upregulation of MHC II genes in through transcriptomic analysis, we next sought to understand whether there was any altered response in CDKL5-mutant iAstros. Both TNFα and IFNγ levels have been reported to be elevated in the blood of CDD patients [92], meaning that the response of CDKL5-mutant astrocytes to elevated levels of these cytokines could be biologically relevant. TNFα exposure elicited a significantly reduced release of both IL-6 and CXCL10 from CDKL5-mutant iAstros compared to WT. A reduced response to TNFα could result in altered glutamate release from CDKL5-null astrocytes. Interestingly, Dumont et al. describe a significant downregulation of the astrocytic glutamate transporter GLAST in WT astrocytes following TNFα exposure, along with an increase in GLT1 expression [93]. Altered glutamate regulation by CDKL5-null astrocytes would likely contribute to a state of neuronal hyperexcitability. In addition, the reduced release of IL-6 by CDKL5-null astrocytes may in itself have epileptogenic functions [94]. IL-6 has been shown to be protective against glutamate excitotoxity, as well as important for neuronal differentiation [95, 96]. Thus, dysregulation of any of these processes through aberrant TNFα response could disrupt network balance and increase seizure susceptibility. Finally, the increased release of CXCL10 in response to IFNγ could have many functional effects on astrocytes and neighbouring cells in the context of CDD. For example, astrocytic CXCL10 has been shown to bind CXCR3 on neurons to initiate neuronal ferroptosis via phosphorylation of STAT3 and enhance seizure susceptibility [97].

Phosphorylation of the known CDKL5 target EB2 was investigated using antibodies for total EB2 protein as well as pEB2 at Ser222. Interestingly, an upregulation of total EB2 protein expression was seen, along with a constant level of pEB2 between WT and CDLK5-mutant iAstros. This resulted in a net decrease of pEB2/EB2 in the CDKL5-mutant iAstros, suggesting that CDKL5 deficiency results in a lower rate of EB2 phosphorylation. The finding that the ratio of pEB2/EB2 is significantly reduced in CDKL5-null astrocytes potentially suggests functional activity of CDKL5 in WT iAstros.

### Organotypic slice culture models of CDD

Given the findings that mouse astrocytes have limited utility in the study of astrocytes in CDD, and the network-relevant dysfunction observed in CDKL5-mutant iAstros, we turned our focus on the generation of a robust *in vitro* human model of CDD, to facilitate future investigations. We therefore utilised shRNA AAV vectors to knockdown CDKL5 in human foetal cortical organotypic slice cultures and to study the functional effect of CDKL5 deficiency in these slices. We also aimed to recapitulate this model in human adult cortical organotypic slice cultures, so as to study the differences in CDKL5 function in the developing vs. mature brain, and thus in developing vs. mature astrocytes. Knockdown of CDKL5 in foetal and adult human cortical tissue had striking similarities; both resulted in trends towards increased spontaneous network activity and increased AQP4 and GFAP expression. However, subtle differences such as the lack of phosphorylated EB2 in the adult tissue hints at a more nuanced functionality of CDKL5 in the developing vs. mature human brain. Given the apparent differences between mouse models of CDD and the human disease phenotype, also reflected in mouse vs. human astrocytes, the generation of novel human models of CDD represents an important step towards understanding molecular mechanisms underlying the disorder.

Synergy between human models presented in this work highlights the value in a combinatory approach for the study of complex neurodevelopmental disorders such as CDD, particularly where representative mouse models do not exist. For example, we show that the water channel AQP4 was upregulated in CDKL5 shRNA adult human organotypic brain slice cultures as well as in CDKL5-mutant iAstros. AQP4 has been shown to be upregulated in TSC astrocytes independent of seizure exposure [98], and dysregulation of AQP4 in CDKL5-null astrocytes could lead to altered osmotic and K+ dynamics and contribute to altered neuronal excitability in CDD. Furthermore, disruption of AQP4 function could alter blood brain barrier permeability. Recent studies have underscored the critical link between dysfunctions in this channel and the pathophysiology of ASD [99]. AQP4 has been further implicated in epilepsy due to its function in maintaining osmotic homeostasis in the brain, and therefore in the control of extracellular space volume and ion/neurotransmitter concentration [100]. For example, AQP4 has been shown to alter spatial potassium buffering and could increase seizure susceptibility through this dysregulation of ion concentration [101]. However, AQP4 has also been linked to cytokine production – offering a potential mechanism through which the upregulation of this water channel may contribute to the altered cytokine response of CDKL5-null astrocytes [102]. Moreover, i*n vitro*, LPS treatment revealed significantly higher cytokine release (but not production) in WT astrocytes compared to AQP4-null astrocytes, highlighting the role of AQP4 in cytokine-mediated response to stimuli [102]. This is further supported in more recent work which showed that AQP4 was involved in the release of proinflammatory cytokines following seizure induction [103]. Therefore, the upregulation of may be closely linked to the basal altered inflammatory gene expression profile and cytokine response seen in CDKL5-mutant iAstros. The fact that this AQP4 upregulation is also demonstrated in the human organotypic slice culture model of CDD suggests this could be conserved in the CDD patient brain. Moreover, an upregulation of the astrogliosis marker GFAP is also noted in the CDKL5 shRNA treated human organotypic slice culture models, further supporting a function of CDKL5 in the mediation of inflammatory phenotypes in astrocytes.

### Species differences underscore the importance of human CDD models

In mouse models of CDD, spontaneous seizure activity has been identified infrequently and only in aged adult mice of some genotypes [104]. One explanation supporting an astrocytic role in the phenotypic differences between species is the apparent increase in CDKL5 expression as mice age. For example, Cdkl5 has been shown to be significantly upregulated in astrocytes in the hypothalamus in aged mice compared to adult mice [105]. Significant upregulation of CDKL5 in the hypothalamus was accompanied by an upregulation of multiple genes relating to inflammation and reactivity in this region [105], potentially reinforcing the link between CDKL5 and inflammatory processes identified in the human iAstros.

Another underlying contributor to species phenotypic differences could lie within astrocytic inflammatory signalling. A key finding in CDKL5-null human astrocytes was the significant upregulation of MHC Class II genes and altered inflammatory response. Critically, this transcriptomic dysregulation was not recapitulated in the CDKL5-null mouse astrocytes. Divergent roles in inflammation have been described in human and mouse astrocytes, and, congruously, the data presented in this paper suggests that CDKL5 may differentially affect immunological processes between the two species. This reflects key differences in cytokine signalling between human and mouse astrocytes reported elsewhere [66]. Moreover, Li et al. reported a stronger response to TNFα in human astrocytes compared to mouse, specifically including induction of antigen-presentation genes in human but not mouse astrocytes [66]. This is particularly interesting given the previous report by Galvani et al. that TNFα release from microglia is increased in CDD [106], and the altered response to TNFa reported in CDKL5-null iAstros. Therefore, it’s possible that CDKL5-null mouse astrocytes do not demonstrate an aberrant response to increased TNFα or other stimulatory cytokines in the way that CDKL5-null human astrocytes were shown to, and this could contribute to the lack of seizures in mouse models. Importantly, the increase of both CDKL5 expression and inflammatory gene expression concurrently in aged mouse astrocytes described above [105], combined with the presence of spontaneous seizures only in aged mice [104], offers direct support for this species difference.

### General conclusions

In this study, we evaluated both mouse and human models of CDD astrocytes. Mouse astrocytes appeared to exhibit lower expression of CDKL5 than human astrocytes and, whilst cytoskeletal transcriptomic dysregulation was present, the loss of expression of the kinase resulted in a lower dysregulation of gene expression in these cells. Meanwhile, key mechanisms of CDKL5 function were dysregulated in human astrocytes including cytoskeletal organisation and inflammatory response. The discrepancy between CDKL5 function in astrocytes between the two species potentially lies with the fact that the affected processes are much less crucial in WT mouse astrocytes than their human counterparts, emphasised by lower expression of CDKL5 in these cells. Species astrocytic differences may contribute to the lack of spontaneous seizures in CDD mouse models. Human models could therefore be utilised for future studies of molecular and network mechanisms in CDD. To facilitate this, proof-of-concept for novel, functional models of CDD utilising CDKL5 shRNA AAV knockdown in foetal and adult human organotypic brain slice cultures was demonstrated. These models further recapitulated expression changes seen in iPSC-derived astrocytes and could be used for future study of both astrocytes and other brain cell-types in CDD. Testing therapeutic interventions in CDD models with network connectivity such as the human organotypic slice cultures presented here, could further validate their therapeutic utility.

## Acknowledgements

Richard Reynolds for critical reading of this manuscript; Simone Di Giovanni and Alessandra Renieri (University of Sienna) for providing iPSC lines; Yunan Gao, Elaine Irvine and Sila Ultanir (Francis Crick Institute) for providing mice; Dirk Grimm (Heidelberg University) for the DJ serotype plasmid. We thank all those involved in donating the foetal tissue samples, as well as the staff of the HDBR tissue bank for their invaluable assistance.

## Author contributions

CP and NDM designed research, performed experiments and wrote initial draft; CJAC performed surgical operations; all other authors contributed to data generation.

## Data availability

All data sets are available on request.

## Ethics

Mice were group housed on a 12/12 dark/light cycle, and food and water were given *ad libitum*. All animal procedures were approved by the local ethical committee and performed in accordance with the United Kingdom Animals Scientific Procedures Act (1986) and associated guidelines.

## Abbreviations

CDKL5: Cyclin-dependent kinase-like 5
CDD: CDKL5 Deficiency disorder
GFAP: glial fibrillary acidic protein
iPSC: inducible Pluripotent stem cells
RNAseq: RNA sequencing
TNFα: Tumour necrosis factor alpha

## References

1. Olson HE, Demarest ST, Pestana-Knight EM, et al (2019) Cyclin-Dependent Kinase-Like 5 Deficiency Disorder: Clinical Review. Pediatr Neurol 97:18–25. 10.1016/j.pediatrneurol.2019.02.015

2. Leonard H, Junaid M, Wong K, et al (2021) Exploring quality of life in individuals with a severe developmental and epileptic encephalopathy, CDKL5 Deficiency Disorder. Epilepsy Res 169:106521. 10.1016/j.eplepsyres.2020.106521

3. Leonard H, Downs J, Benke TA, et al (2022) CDKL5 deficiency disorder: clinical features, diagnosis, and management. Lancet Neurol 21:563–576. 10.1016/S1474-4422(22)00035-7

4. Hong W, Haviland I, Pestana-Knight E, et al (2022) CDKL5 Deficiency Disorder-Related Epilepsy: A Review of Current and Emerging Treatment. CNS Drugs 36:591–604. 10.1007/s40263-022-00921-5

5. Fehr S, Downs J, Ho G, et al (2016) Functional abilities in children and adults with the CDKL5 disorder. Am J Med Genet A 170:2860–2869. 10.1002/ajmg.a.37851

6. Jakimiec M, Paprocka J, Śmigiel R (2020) CDKL5 Deficiency Disorder—A Complex Epileptic Encephalopathy. Brain Sci 10:107. 10.3390/brainsci10020107

7. Siri B, Varesio C, Freri E, et al (2021) CDKL5 deficiency disorder in males: Five new variants and review of the literature. European Journal of Paediatric Neurology 33:9–20. 10.1016/j.ejpn.2021.04.007

8. Knight EMP, Amin S, Bahi-Buisson N, et al (2022) Safety and efficacy of ganaxolone in patients with CDKL5 deficiency disorder: results from the double-blind phase of a randomised, placebo-controlled, phase 3 trial. Lancet Neurol 21:417–427. 10.1016/S1474-4422(22)00077-1

9. Montini E, Andolfi G, Caruso A, et al (1998) Identification and Characterization of a Novel Serine– Threonine Kinase Gene from the Xp22 Region. Genomics 51:427–433. 10.1006/geno.1998.5391

10. Hector RD, Dando O, Landsberger N, et al (2016) Characterisation of CDKL5 Transcript Isoforms in Human and Mouse. PLoS One 11:e0157758. 10.1371/journal.pone.0157758

11. Demarest ST, Olson HE, Moss A, et al (2019) CDKL5 deficiency disorder: Relationship between genotype, epilepsy, cortical visual impairment, and development. Epilepsia 60:1733–1742. 10.1111/epi.16285

12. Rusconi L, Salvatoni L, Giudici L, et al (2008) CDKL5 Expression Is Modulated during Neuronal Development and Its Subcellular Distribution Is Tightly Regulated by the C-terminal Tail. Journal of Biological Chemistry 283:30101–30111. 10.1074/jbc.M804613200

13. Terzic B, Davatolhagh MF, Ho Y, et al (2021) Temporal manipulation of Cdkl5 reveals essential postdevelopmental functions and reversible CDKL5 deficiency disorder–related deficits. Journal of Clinical Investigation 131:. 10.1172/JCI143655

14. Van Bergen NJ, Massey S, Quigley A, et al (2022) CDKL5 deficiency disorder: molecular insights and mechanisms of pathogenicity to fast-track therapeutic development. Biochem Soc Trans 50:1207–1224. 10.1042/BST20220791

15. Muñoz IM, Morgan ME, Peltier J, et al (2018) Phosphoproteomic screening identifies physiological substrates of the CDKL5 kinase. EMBO J 37:. 10.15252/embj.201899559

16. Amendola E, Zhan Y, Mattucci C, et al (2014) Mapping Pathological Phenotypes in a Mouse Model of CDKL5 Disorder. PLoS One 9:e91613. 10.1371/journal.pone.0091613

17. Zhu Y-C, Li D, Wang L, et al (2013) Palmitoylation-dependent CDKL5–PSD-95 interaction regulates synaptic targeting of CDKL5 and dendritic spine development. Proceedings of the National Academy of Sciences 110:9118–9123. 10.1073/pnas.1300003110

18. Della Sala G, Putignano E, Chelini G, et al (2016) Dendritic Spine Instability in a Mouse Model of CDKL5 Disorder Is Rescued by Insulin-like Growth Factor 1. Biol Psychiatry 80:302–311. 10.1016/j.biopsych.2015.08.028

19. Baltussen LL, Negraes PD, Silvestre M, et al (2018) Chemical genetic identification of CDKL5 substrates reveals its role in neuronal microtubule dynamics. EMBO J 37:. 10.15252/embj.201899763

20. Sampedro-Castañeda M, Baltussen LL, Lopes AT, et al (2023) Epilepsy-linked kinase CDKL5 phosphorylates voltage-gated calcium channel Cav2.3, altering inactivation kinetics and neuronal excitability. Nat Commun 14:7830. 10.1038/s41467-023-43475-w

21. Simões de Oliveira L, O’Leary HE, Nawaz S, et al (2024) Enhanced hippocampal LTP but normal NMDA receptor and AMPA receptor function in a rat model of CDKL5 deficiency disorder. Mol Autism 15:28. 10.1186/s13229-024-00601-9

22. Varela T, Varela D, Martins G, et al (2022) Cdkl5 mutant zebrafish shows skeletal and neuronal alterations mimicking human CDKL5 deficiency disorder. Sci Rep 12:9325. 10.1038/s41598-022-13364-1

23. Serrano RJ, Lee C, Douek AM, et al (2022) Novel preclinical model for CDKL5 deficiency disorder. Dis Model Mech 15:. 10.1242/dmm.049094

24. Wang I-TJ, Allen M, Goffin D, et al (2012) Loss of CDKL5 disrupts kinome profile and event-related potentials leading to autistic-like phenotypes in mice. Proceedings of the National Academy of Sciences 109:21516–21521. 10.1073/pnas.1216988110

25. Tang S, Wang I-TJ, Yue C, et al (2017) Loss of CDKL5 in Glutamatergic Neurons Disrupts Hippocampal Microcircuitry and Leads to Memory Impairment in Mice. The Journal of Neuroscience 37:7420–7437. 10.1523/JNEUROSCI.0539-17.2017

26. Wang H, Zhu Z, Li Y, et al (2021) CDKL5 deficiency in forebrain glutamatergic neurons results in recurrent spontaneous seizures. Epilepsia 62:517–528. 10.1111/epi.16805

27. Medici G, Tassinari M, Galvani G, et al (2022) Expression of a Secretable, Cell-Penetrating CDKL5 Protein Enhances the Efficacy of Gene Therapy for CDKL5 Deficiency Disorder. Neurotherapeutics 19:1886–1904. 10.1007/s13311-022-01295-8

28. Gao Y, Irvine EE, Eleftheriadou I, et al (2020) Gene replacement ameliorates deficits in mouse and human models of cyclin-dependent kinase-like 5 disorder. Brain 143:811–832. 10.1093/brain/awaa028

29. Baldwin KT, Eroglu C (2017) Molecular mechanisms of astrocyte-induced synaptogenesis. Curr Opin Neurobiol 45:113–120. 10.1016/j.conb.2017.05.006

30. Christopherson KS, Ullian EM, Stokes CCA, et al (2005) Thrombospondins Are Astrocyte-Secreted Proteins that Promote CNS Synaptogenesis. Cell 120:421–433. 10.1016/j.cell.2004.12.020

31. Siracusa R, Fusco R, Cuzzocrea S (2019) Astrocytes: Role and Functions in Brain Pathologies. Front Pharmacol 10:. 10.3389/fphar.2019.01114

32. von Bartheld CS, Bahney J, Herculano-Houzel S (2016) The search for true numbers of neurons and glial cells in the human brain: A review of 150 years of cell counting. Journal of Comparative Neurology 524:3865–3895. 10.1002/cne.24040

33. Coulter DA, Steinhauser C (2015) Role of Astrocytes in Epilepsy. Cold Spring Harb Perspect Med 5:a022434–a022434. 10.1101/cshperspect.a022434

34. Oberheim NA, Tian G-F, Han X, et al (2008) Loss of Astrocytic Domain Organization in the Epileptic Brain. The Journal of Neuroscience 28:3264–3276. 10.1523/JNEUROSCI.4980-07.2008

35. Bedner P, Dupper A, Hüttmann K, et al (2015) Astrocyte uncoupling as a cause of human temporal lobe epilepsy. Brain 138:1208–1222. 10.1093/brain/awv067

36. Schröder W, Hinterkeuser S, Seifert G, et al (2000) Functional and Molecular Properties of Human Astrocytes in Acute Hippocampal Slices Obtained from Patients with Temporal Lobe Epilepsy. Epilepsia 41:. 10.1111/j.1528-1157.2000.tb01578.x

37. Evans JC, Archer HL, Colley JP, et al (2005) Early onset seizures and Rett-like features associated with mutations in CDKL5. European Journal of Human Genetics 13:1113–1120. 10.1038/sj.ejhg.5201451

38. Swartzlander DB, Propson NE, Roy ER, et al (2018) Concurrent cell type–specific isolation and profiling of mouse brains in inflammation and Alzheimer’s disease. JCI Insight 3:. 10.1172/jci.insight.121109

39. Fakhr E, Zare F, Teimoori-Toolabi L (2016) Precise and efficient siRNA design: a key point in competent gene silencing. Cancer Gene Ther 23:73–82. 10.1038/cgt.2016.4

40. Brummelkamp TR, Bernards R, Agami R (2002) A System for Stable Expression of Short Interfering RNAs in Mammalian Cells. Science (1979) 296:550–553. 10.1126/science.1068999

41. Mcintyre GJ, Yu Y-H, Lomas M, Fanning GC (2011) The effects of stem length and core placement on shRNA activity. BMC Mol Biol 12:34. 10.1186/1471-2199-12-34

42. McLeod F, Dimtsi A, Marshall AC, et al (2023) Altered synaptic connectivity in an in vitro human model of STXBP1 encephalopathy. Brain 146:850–857. 10.1093/brain/awac396

43. Stael S, Miller LP, Fernández-Fernández ÁD, Van Breusegem F (2022) Detection of Damage-Activated Metacaspase Activity by Western Blot in Plants. pp 127–137

44. Livak KJ, Schmittgen TD (2001) Analysis of Relative Gene Expression Data Using Real-Time Quantitative PCR and the 2−ΔΔCT Method. Methods 25:402–408. 10.1006/meth.2001.1262

45. Andrews S (2010) FastQC: A Quality Control Tool for High Throughput Sequence Data [Online]. In: http://www.bioinformatics.babraham.ac.uk/projects/fastqc/

46. Chen S, Zhou Y, Chen Y, Gu J (2018) fastp: an ultra-fast all-in-one FASTQ preprocessor. Bioinformatics 34:i884–i890. 10.1093/bioinformatics/bty560

47. Patro R, Duggal G, Love MI, et al (2017) Salmon provides fast and bias-aware quantification of transcript expression. Nat Methods 14:417–419. 10.1038/nmeth.4197

48. Soneson C, Love MI, Robinson MD (2015) Differential analyses for RNA-seq: transcript-level estimates improve gene-level inferences. F1000Res 4:1521. 10.12688/f1000research.7563.1

49. Tarazona S, Furió-Tarí P, Turrà D, et al (2015) Data quality aware analysis of differential expression in RNA-seq with NOISeq R/Bioc package. Nucleic Acids Res gkv711. 10.1093/nar/gkv711

50. Yu G, Wang L-G, Han Y, He Q-Y (2012) clusterProfiler: an R Package for Comparing Biological Themes Among Gene Clusters. OMICS 16:284–287. 10.1089/omi.2011.0118

51. Szklarczyk D, Kirsch R, Koutrouli M, et al (2023) The STRING database in 2023: protein–protein association networks and functional enrichment analyses for any sequenced genome of interest. Nucleic Acids Res 51:D638–D646. 10.1093/nar/gkac1000

52. Love MI, Huber W, Anders S (2014) Moderated estimation of fold change and dispersion for RNA-seq data with DESeq2. Genome Biol 15:550. 10.1186/s13059-014-0550-8

53. Stephens M, Carbonetto P, Gerard D, et al (2023) ashr: Methods for Adaptive Shrinkage, using Empirical Bayes

54. Blighe K, Rana S, Lewis M (2024) Publication-ready volcano plots with enhanced colouring and labeling. version 1.24.0

55. Yu G (2024) enrichplot: Visualization of Functional Enrichment Result

56. Kolde R (2019) pheatmap: Pretty Heatmaps. CRAN: Contributed Packages

57. TCW J, Wang M, Pimenova AA, et al (2017) An Efficient Platform for Astrocyte Differentiation from Human Induced Pluripotent Stem Cells. Stem Cell Reports 9:600–614. 10.1016/j.stemcr.2017.06.018

58. Leventoux N, Morimoto S, Imaizumi K, et al (2020) Human Astrocytes Model Derived from Induced Pluripotent Stem Cells. Cells 9:2680. 10.3390/cells9122680

59. Hedegaard A, Monzón-Sandoval J, Newey SE, et al (2020) Pro-maturational Effects of Human iPSC-Derived Cortical Astrocytes upon iPSC-Derived Cortical Neurons. Stem Cell Reports 15:38–51. 10.1016/j.stemcr.2020.05.003

60. Lorenzi L, Chiu H-S, Avila Cobos F, et al (2021) The RNA Atlas expands the catalog of human non-coding RNAs. Nat Biotechnol 39:1453–1465. 10.1038/s41587-021-00936-1

61. Voss AJ, Lanjewar SN, Sampson MM, et al (2023) Identification of ligand–receptor pairs that drive human astrocyte development. Nat Neurosci 26:1339–1351. 10.1038/s41593-023-01375-8

62. Zhang Y, Chen K, Sloan SA, et al (2014) An RNA-Sequencing Transcriptome and Splicing Database of Glia, Neurons, and Vascular Cells of the Cerebral Cortex. The Journal of Neuroscience 34:11929–11947. 10.1523/JNEUROSCI.1860-14.2014

63. Zhang Y, Sloan SA, Clarke LE, et al (2016) Purification and Characterization of Progenitor and Mature Human Astrocytes Reveals Transcriptional and Functional Differences with Mouse. Neuron 89:37–53. 10.1016/j.neuron.2015.11.013

64. Ma N-X, Yin J-C, Chen G (2019) Transcriptome Analysis of Small Molecule–Mediated Astrocyte-to-Neuron Reprogramming. Front Cell Dev Biol 7:. 10.3389/fcell.2019.00082

65. Crowe EP, Tuzer F, Gregory BD, et al (2016) Changes in the Transcriptome of Human Astrocytes Accompanying Oxidative Stress-Induced Senescence. Front Aging Neurosci 8:. 10.3389/fnagi.2016.00208

66. Li J, Pan L, Pembroke WG, et al (2021) Conservation and divergence of vulnerability and responses to stressors between human and mouse astrocytes. Nat Commun 12:3958. 10.1038/s41467-021-24232-3

67. Cheng Y-T, Luna-Figueroa E, Woo J, et al (2023) Inhibitory input directs astrocyte morphogenesis through glial GABABR. Nature 617:369–376. 10.1038/s41586-023-06010-x

68. Lattke M, Goldstone R, Ellis JK, et al (2021) Extensive transcriptional and chromatin changes underlie astrocyte maturation in vivo and in culture. Nat Commun 12:4335. 10.1038/s41467-021-24624-5

69. Erickson EK, Blednov YA, Harris RA, Mayfield RD (2019) Glial gene networks associated with alcohol dependence. Sci Rep 9:10949. 10.1038/s41598-019-47454-4

70. Roboon J, Hattori T, Nguyen DT, et al (2022) Isolation of ferret astrocytes reveals their morphological, transcriptional, and functional differences from mouse astrocytes. Front Cell Neurosci 16:. 10.3389/fncel.2022.877131

71. Schulte A, Bieniussa L, Gupta R, et al (2022) Homeostatic calcium fluxes, ER calcium release, SOCE, and calcium oscillations in cultured astrocytes are interlinked by a small calcium toolkit. Cell Calcium 101:102515. 10.1016/j.ceca.2021.102515

72. Schmid RS, Simon JM, Vitucci M, et al (2016) Core pathway mutations induce de-differentiation of murine astrocytes into glioblastoma stem cells that are sensitive to radiation but resistant to temozolomide. Neuro Oncol 18:962–973. 10.1093/neuonc/nov321

73. Yao Z, van Velthoven CTJ, Nguyen TN, et al (2021) A taxonomy of transcriptomic cell types across the isocortex and hippocampal formation. Cell 184:3222–3241.e26. 10.1016/j.cell.2021.04.021

74. Yao Z, Liu H, Xie F, et al (2021) A transcriptomic and epigenomic cell atlas of the mouse primary motor cortex. Nature 598:103–110. 10.1038/s41586-021-03500-8

75. Steuernagel L, Lam BYH, Klemm P, et al (2022) HypoMap—a unified single-cell gene expression atlas of the murine hypothalamus. Nat Metab 4:1402–1419. 10.1038/s42255-022-00657-y

76. Batiuk MY, Martirosyan A, Wahis J, et al (2020) Identification of region-specific astrocyte subtypes at single cell resolution. Nat Commun 11:1220. 10.1038/s41467-019-14198-8

77. Hochgerner H, Zeisel A, Lönnerberg P, Linnarsson S (2018) Conserved properties of dentate gyrus neurogenesis across postnatal development revealed by single-cell RNA sequencing. Nat Neurosci 21:290–299. 10.1038/s41593-017-0056-2

78. Siletti K, Hodge R, Mossi Albiach A, et al (2023) Transcriptomic diversity of cell types across the adult human brain. Science (1979) 382:. 10.1126/science.add7046

79. Jorstad NL, Close J, Johansen N, et al (2023) Transcriptomic cytoarchitecture reveals principles of human neocortex organization. Science (1979) 382:. 10.1126/science.adf6812

80. Seeker LA, Bestard-Cuche N, Jäkel S, et al (2023) Brain matters: unveiling the distinct contributions of region, age, and sex to glia diversity and CNS function. Acta Neuropathol Commun 11:84. 10.1186/s40478-023-01568-z

81. Jäkel S, Agirre E, Mendanha Falcão A, et al (2019) Altered human oligodendrocyte heterogeneity in multiple sclerosis. Nature 566:543–547. 10.1038/s41586-019-0903-2

82. Phan BN, Ray MH, Xue X, et al (2024) Single nuclei transcriptomics in human and non-human primate striatum in opioid use disorder. Nat Commun 15:878. 10.1038/s41467-024-45165-7

83. Darmanis S, Sloan SA, Zhang Y, et al (2015) A survey of human brain transcriptome diversity at the single cell level. Proceedings of the National Academy of Sciences 112:7285–7290. 10.1073/pnas.1507125112

84. Lindsay SJ, Xu Y, Lisgo SN, et al (2016) HDBR Expression: A Unique Resource for Global and Individual Gene Expression Studies during Early Human Brain Development. Front Neuroanat 10:. 10.3389/fnana.2016.00086

85. McLeod F, Marzo A, Podpolny M, et al (2017) Evaluation of Synapse Density in Hippocampal Rodent Brain Slices. Journal of Visualized Experiments. 10.3791/56153-v

86. Silvestre M, Dempster K, Mihaylov SR, et al (2024) Cell type-specific expression, regulation and compensation of CDKL5 activity in mouse brain. Mol Psychiatry 29:1844–1856. 10.1038/s41380-024-02434-7

87. Stüve O, Youssef S, Slavin AJ, et al (2002) The Role of the MHC Class II Transactivator in Class II Expression and Antigen Presentation by Astrocytes and in Susceptibility to Central Nervous System Autoimmune Disease. The Journal of Immunology 169:6720–6732. 10.4049/jimmunol.169.12.6720

88. Greer SF, Harton JA, Linhoff MW, et al (2004) Serine Residues 286, 288, and 293 within the CIITA: A Mechanism for Down-Regulating CIITA Activity through Phosphorylation. The Journal of Immunology 173:376–383. 10.4049/jimmunol.173.1.376

89. Morgan JE, Shanderson RL, Boyd NH, et al (2015) The class II transactivator (CIITA) is regulated by post-translational modification cross-talk between ERK1/2 phosphorylation, mono-ubiquitination and Lys63 ubiquitination. Biosci Rep 35:. 10.1042/BSR20150091

90. Voong LN, Slater AR, Kratovac S, Cressman DE (2008) Mitogen-activated protein kinase ERK1/2 regulates the class II transactivator. J Biol Chem 283:9031–9. 10.1074/jbc.M706487200

91. Loi M, Trazzi S, Fuchs C, et al (2020) Increased DNA Damage and Apoptosis in CDKL5-Deficient Neurons. Mol Neurobiol 57:2244–2262. 10.1007/s12035-020-01884-8

92. Cortelazzo A, de Felice C, Leoncini S, et al (2017) Inflammatory protein response in CDKL5-Rett syndrome: evidence of a subclinical smouldering inflammation. Inflammation Research 66:269–280. 10.1007/s00011-016-1014-2

93. Dumont AO, Goursaud S, Desmet N, Hermans E (2014) Differential Regulation of Glutamate Transporter Subtypes by Pro-Inflammatory Cytokine TNF-α in Cortical Astrocytes from a Rat Model of Amyotrophic Lateral Sclerosis. PLoS One 9:e97649. 10.1371/journal.pone.0097649

94. Li G, Bauer S, Nowak M, et al (2011) Cytokines and epilepsy. Seizure 20:249–256. 10.1016/j.seizure.2010.12.005

95. Oh J, McCloskey MA, Blong CC, et al (2010) Astrocyte-derived interleukin-6 promotes specific neuronal differentiation of neural progenitor cells from adult hippocampus. J Neurosci Res 88:2798–2809. 10.1002/jnr.22447

96. Peng Y-P, Qiu Y-H, Lu J-H, Wang J-J (2005) Interleukin-6 protects cultured cerebellar granule neurons against glutamate-induced neurotoxicity. Neurosci Lett 374:192–196. 10.1016/j.neulet.2004.10.069

97. Liang P, Zhang X, Zhang Y, et al (2023) Neurotoxic A1 astrocytes promote neuronal ferroptosis via CXCL10/CXCR3 axis in epilepsy. Free Radic Biol Med 195:329–342. 10.1016/j.freeradbiomed.2023.01.002

98. Short B, Kozek L, Harmsen H, et al (2019) Cerebral aquaporin-4 expression is independent of seizures in tuberous sclerosis complex. Neurobiol Dis 129:93–101. 10.1016/j.nbd.2019.05.003

99. Abbasian V, Davoudi S, Vahabzadeh A, et al (2025) Astroglial Kir4.1 and AQP4 Channels: Key Regulators of Potassium Homeostasis and Their Implications in Autism Spectrum Disorders. Cell Mol Neurobiol 45:56. 10.1007/s10571-025-01574-w

100. Verhoog QP, Holtman L, Aronica E, van Vliet EA (2020) Astrocytes as Guardians of Neuronal Excitability: Mechanisms Underlying Epileptogenesis. Front Neurol 11:. 10.3389/fneur.2020.591690

101. Bonosi L, Benigno UE, Musso S, et al (2023) The Role of Aquaporins in Epileptogenesis—A Systematic Review. Int J Mol Sci 24:11923. 10.3390/ijms241511923

102. Li L, Zhang H, Varrin-Doyer M, et al (2011) Proinflammatory role of aquaporin-4 in autoimmune neuroinflammation. The FASEB Journal 25:1556–1566. 10.1096/fj.10-177279

103. Lei S, He Y, Zhu Z, et al (2020) Inhibition of NMDA Receptors Downregulates Astrocytic AQP4 to Suppress Seizures. Cell Mol Neurobiol 40:1283–1295. 10.1007/s10571-020-00813-6

104. Mulcahey PJ, Tang S, Takano H, et al (2020) Aged heterozygous Cdkl5 mutant mice exhibit spontaneous epileptic spasms. Exp Neurol 332:113388. 10.1016/j.expneurol.2020.113388

105. Boisvert MM, Erikson GA, Shokhirev MN, Allen NJ (2018) The Aging Astrocyte Transcriptome from Multiple Regions of the Mouse Brain. Cell Rep 22:269–285. 10.1016/j.celrep.2017.12.039

106. Galvani G, Mottolese N, Gennaccaro L, et al (2021) Inhibition of microglia overactivation restores neuronal survival in a mouse model of CDKL5 deficiency disorder. J Neuroinflammation 18:155. 10.1186/s12974-021-02204-0

107. Yao Z, van Velthoven CTJ, Nguyen TN, et al (2021) A taxonomy of transcriptomic cell types across the isocortex and hippocampal formation. Cell 184:3222–3241.e26. 10.1016/j.cell.2021.04.021

108. Yao Z, Liu H, Xie F, et al (2021) A transcriptomic and epigenomic cell atlas of the mouse primary motor cortex. Nature 598:103–110. 10.1038/s41586-021-03500-8

