## Supplemental data for "Astroglial Dysfunction in Models of CDKL5 Deficiency Disorder" (from another version of this preprint: version 1, 2026-06-10)

### SUPPLEMENTARY INFORMATION

#### Supplemental Tables

| TARGET | HOST | SOURCE | CLONALITY | DILUTION | | |
|---|---|---|---|---|---|---|
| | | | | WB | ICC | IHC |
| CDKL5 | Mouse | Santa Cruz, Sc-376314 | Monoclonal | 1:500-1:1000 | - | - |
| ALDH1L1 | Rabbit | Abcam, ab190293 | Polyclonal | - | 1:500 | - |
| AQP4 | Rabbit | ProteinTech, 16473-AP | Polyclonal | 1:1000 - 1:5000 | 1:100 | - |
| AQP4 | Rabbit | Alomone, 249-323 | Polyclonal | - | 1:2000 | 1:1000 |
| GFAP | Rabbit | Abcam, ab68428 | Polyclonal | 1:1000 - 1:4000 | 1:250 | - |
| GFAP | Chicken | Abcam, ab4674 | Polyclonal | - | - | 1:2000 |
| GFP | Rabbit | Abcam, ab290 | Polyclonal | - | - | 1:1000 |
| NeuN | Mouse | Abcam, ab104224 | Monoclonal | - | - | 1:1000 |
| GAPDH | Rabbit | Cell Signalling, 2118 | Monoclonal | 1:2500 | - | - |
| pEB2 | Rabbit | Produced and provided by Dr. Sila Ultanir. | Polyclonal | 1:500 - 1:2000 | - | - |
| EB2 | Rat | Abcam, ab45767 | Monoclonal | 1:500 – 1:1000 | - | - |
| S100β | Rabbit | Abcam, Ab52642 | Polyclonal | - | 1:500 | |
| CD44 | Rabbit | Abcam, Ab243894 | Monoclonal | - | 1:500 | - |

Supplemental Table 1 – List of primary antibodies

| NAME | SOURCE | DILUTION | | |
|---|---|---|---|---|
| | | WB | ICC | IHC |
| Donkey anti-Rabbit IgG H&L (HRP) | Abcam, ab16284 | 1:2000 | - | - |
| Goat anti-Mouse IgG (H+L), Alexa Fluor™ 594 | Invitrogen, A11032 | - | 1:1000 | - |
| Goat anti-Rabbit IgG (H+L) Alexa Fluor™ 488 | Invitrogen, A11034 | - | 1:1000 | - |
| Peroxidase AffiniPure™ Goat anti-Mouse HRP | Jackson Laboratories, 115-035-003 | 1:10,000 | - | - |
| Donkey anti-Rabbit IgG (H+L) Alexa Fluor™ 488 | Invitrogen, A21206 | - | - | 1:500 |
| Donkey anti-Mouse IgG (H+L) Alexa Fluor™ 568 | Invitrogen, A10037 | - | - | 1:500 |
| Alexa Fluor® 647 AffiniPure™ Donkey Anti-Chicken IgY (IgG) (H+L) | Jackson Laboratories, 703-605-155 | | | 1:500 |
| Goat anti-Rat IgG (H+L), HRP | Invitrogen, 31470 | 1:5000 | - | - |

Supplemental Table 2 – List of Secondary Antibodies

| PRIMARY AUTHOR | DESCRIPTION | NUMBER OF SAMPLES | REFERENCE |
|---|---|---|---|
| Lorenzi | Cell line (Sciencell) | 3 | [60] |
| Voss | Isolated from foetal cortex | 2 | [61] |
| Zhang | Isolated from foetal cortex | 5 | [63] |
| Zhang | Isolated from adolescent temporal lobe | 2 | [63] |
| Zhang | Isolated from adult temporal lobe | 2 | [63] |
| Ma | Cell line (Sciencell) | 6 | [64] |
| Crowe | Cell line (Sciencell) | 2 | [65] |
| Li | Isolated from foetal cortex | 2 | [66] |

Supplemental Table 3a – List of human bulk RNAseq datasets

| PRIMARY AUTHOR | REGION | AGE CATEGORY | NUMBER OF SAMPLES | REFERENCE |
|---|---|---|---|---|
| Lattke | Cortex | Juvenile | 3 | [68] |
| Lattke | Cortex | Young Adult | 3 | [68] |
| Erickson | Cortex | Young Adult | 17 | [69] |
| Cheng | Hippocampus | Juvenile | 9 | [67] |
| Cheng | Cortex | Juvenile | 4 | [67] |
| Roboon | Cortex | Juvenile | 4 | [70] |
| Schulte | Cortex | Juvenile | 4 | [71] |
| Schmid | Cortex | Juvenile | 2 | [72] |

Supplemental Table 3b – list of mouse bulk RNAseq datasets

| PRIMARY AUTHOR | REGION | SEQUENCING METHOD | AGE | SPECIES | NUMBER OF CELLS | REFERENCE |
|---|---|---|---|---|---|---|
| Yao | Isocortex and hippocampus | scRNAseq | P50-121 | Mouse | 1638333 | [107] |
| Yao | Primary motor cortex | scRNAseq and snRNAseq | P52-59 | Mouse | 18905 | [108] |
| Steuernagel | Hypothalamus | snRNAseq | Various | Mouse | 59785 | [75] |
| Batiuk | Hippocampus | scRNAseq | P53 | Mouse | 820 | [76] |
| Batiuk | Cortex | scRNAseq | P53 | Mouse | 991 | [76] |
| Hochgerner | Hippocampus | scRNAseq | P12-35 | Mouse | 5454 | [77] |
| Siletti | Right Hemisphere | snRNAseq | 29 – 60 years | Human | 158449 | [78] |
| Jorstad | Cortex | scRNAseq | 24 - 57 years | Human | 20759 | [79] |
| Seeker | Primary motor cortex and cerebellum | snRNAseq | 30 – 75 years | Human | 768 | [80] |
| Jakel | White matter | snRNAseq | 35 – 82 years | Human | 392 | [81] |
| Phan | Caudate and Putamen | snRNAseq | 23 – 60 years | Human | 4195 | [82] |
| Darmanis | Cortex (anterior temporal lobe) | scRNAseq | 21 – 63 years | Human | 466 | [83] |

Supplemental Table 3c – List of sc/snRNAseq datsets

| NAME | 5' TO 3' SEQUENCE |
|---|---|
| CDKL5 shRNA | GATCCCCGGG ACAAAGTACC TCAAGTCAAA TTCAAGAGAT TTGACTTGAG GTACTTTGTC CCGGTTTTTT ACGCGTG |
| Scrambled shRNA | GATCCGAACA ATCCCAGGAG ACGCACTAAT TTCAAGAGAA TTAGTGCGTC TCCTGGGATT GTTCTTTTTT ACGCGTG |

Supplemental Table 4 – 5' to 3' shRNA construct cloned into AAV transfer plasmid. Green = sticky ends; orange = siRNA target sequence and reverse complement; red = loop sequence; blue = termination signal.

| Case Number | Age (pcw) | Brain Region | Experiment |
|---|---|---|---|
| FT20 | 17 | Occipital Ctx. | Immunostaining |
| FT21 | 17 | Occipital Ctx. | Immunostaining |
| FT24 | 17 | Occipital Ctx. | Immunostaining |
| FT32 | 17 | Occipital Ctx. | Transduced with shRNA/Scr AAV, MEA, protein and RNA analysis |
| FT27 | 18 | Occipital Ctx. | Transduced with shRNA/Scr AAV, protein and RNA analysis |
| FT36 | 18 | Frontal Ctx | Transduced with shRNA/Scr AAV, protein and RNA analysis |
| FT37 | 16 | Occipital Ctx | Transduced with shRNA/Scr AAV, RNA analysis |

Supplemental Table 5 – Age and region of foetal brain tissue utilised in the current study.

| Case Number | Age (yrs) | Brain Region | Sex | Experiment |
|---|---|---|---|---|
| CE94 | 71 | Right temporal cortex | Male | Transduced with shRNA/Scr AAV. Immunostaining, protein analysis. |
| CE95 | 60 | Left temporal cortex | Female | Transduced with shRNA/Scr AAV, MEA, protein and RNA analysis |
| CE97 | 56 | Right temporal cortex | Male | Transduced with shRNA/Scr AAV, MEA, protein and RNA analysis |
| CE98 | 30 | Inferior temporal gyrus | Female | Transduced with shRNA/Scr AAV, Protein analysis |

Supplemental Table 6 – Age and region of adult cortical brain tissue utilised in the current study.

**Supplemental Figures**

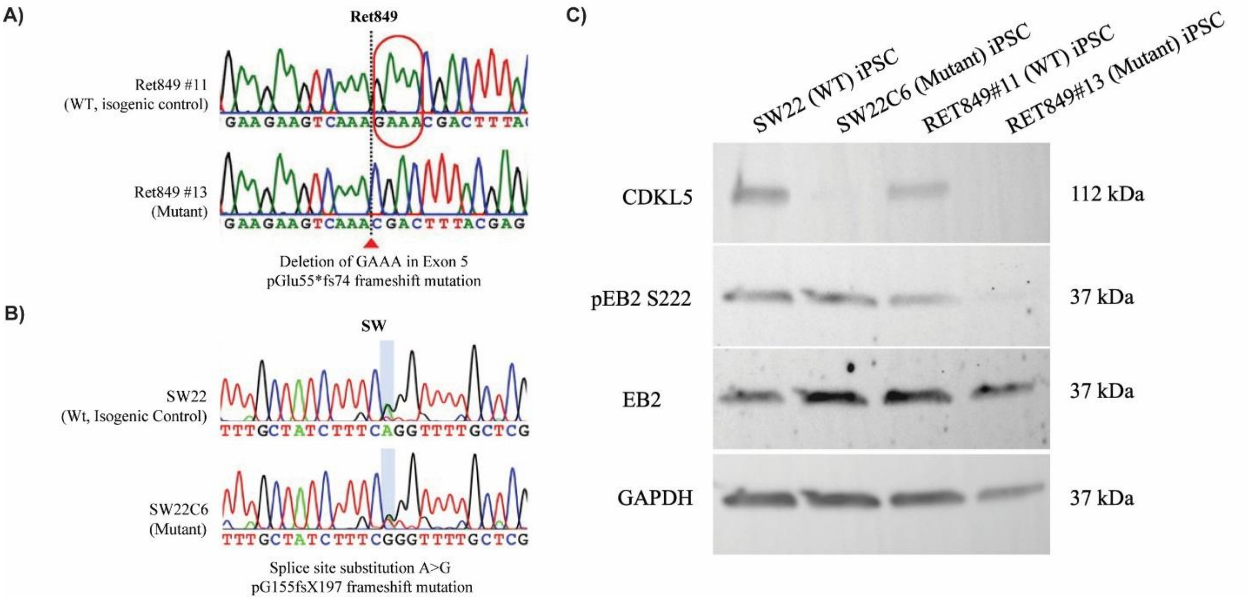


### Supplemental Figure 1: CDKL5 gene variants in iPSCs

**A)** Sequencing of RT-PCR products generated using RNA extracted from Ret849 #11 and #13 iPSCs and located in exon 5 of *CDKL5*. Ret849 #13 has a deletion of GAAA (represented by red circle in Ret849 #11), resulting in a pGlu55*fs74 frameshift mutation. Also appears in Gao et al. 2020 [28]. **B)** Sequencing of PCR product of genomic *CDKL5*, generated from DNA extracted from SW22 and SW22C6 iPSCs. SW22C6 has a deletion at the splice acceptor site for exon 8, highlighted by red line and resulting in a pG155fsX197 frameshift mutation. The A>G substitution at position 298 (B highlighted) in the mutant amplicon was confirmed using different base calling software at AZENTA Life Sciences. **C)** Diminished CDKL5 protein expression in SW22C6 and Ret849 #13 iPSC cell lines was confirmed via Western Blot. Protein from WT and CDKL5 mutant iPSCs was probed for CDKL5, phosphorylated EB2 (pEB2) at Serine 222 (S222), EB2 and GAPDH as a loading control.

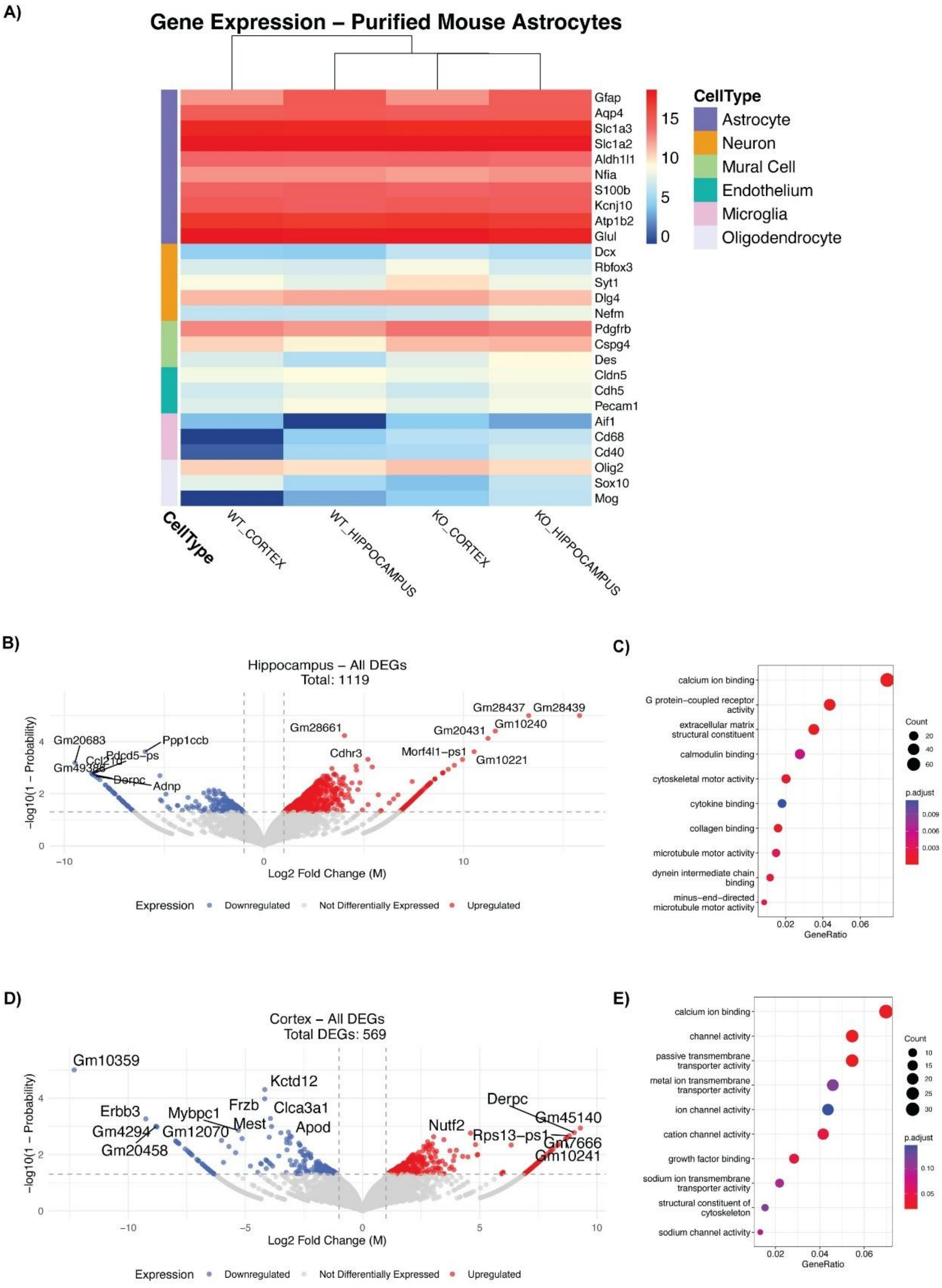


**Supplemental Figure 2: RNAseq Analysis of Astrocytes purified from Cdkl5$^{-/y}$ KO and WT littermate mice.**

**A)** Heatmap showing $\log_2$ normalised counts of genes associated with astrocytes, mural cells, endothelium, neurons, microglia and oligodendrocytes in purified Cdkl5-null (KO) and wildtype (WT) hippocampal and cortical astrocytes. Plot generated in RStudio using pheatmaps package with RColorBrewer.

**B)** Volcano plot showing -log10(1-probability) against $\log_2$ fold change (lfc) differentially expressed genes in KO mouse hippocampal astrocytes compared to WT. Each dot represents a gene, coloured dots represent significantly differentially expressed genes, up- or downregulated according to the legend below each plot. Lfc threshold > 1, probability threshold > 0.95. Dashed line indicates location of threshold.

**C)** Top 10 most significantly enriched Molecular Function Gene Ontology annotations for mouse hippocampus DEGs.

**D)** As for B but Cortex.

**E)** As for C but for Cortex.

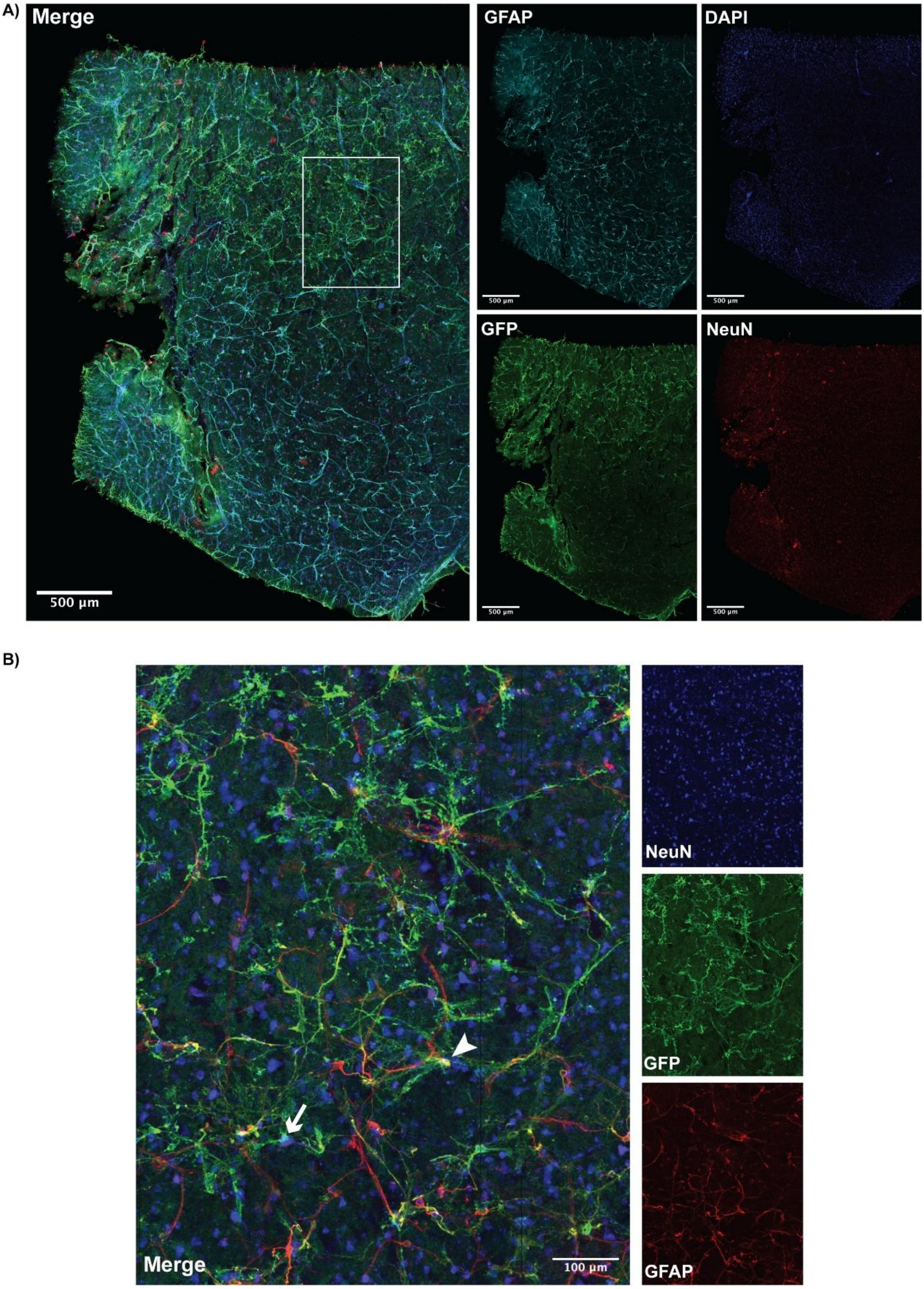


**Supplemental Figure 3: GFP Expression in adult human organotypic slice transduced with Scrambled shRNA.**

**A)** Representative confocal scanning images of human adult cortical slice transduced with AAVDJ-U6-ScrambledshRNA-CMV-eGFP (Scrambled shRNA), and stained with GFAP (cyan), GFP (green), NeuN (red) and the nuclear marker DAPI (blue). Scale bars represent 500 µm. Images were taken using a Zeiss LSM800 Airyscan confocal microscope.
**B)** Zoomed in area of A), position indicated by white box in A). In this image, DAPI is removed for easier visualisation: NeuN is blue, GFP is still green and GFAP is red. Scale bar represents 100 µm. Arrow shows GFP colocalization with NeuN (cyan, neuronal transduction) whilst pointed triangle shows GFP colocalization with GFAP (yellow, astrocytic transduction).

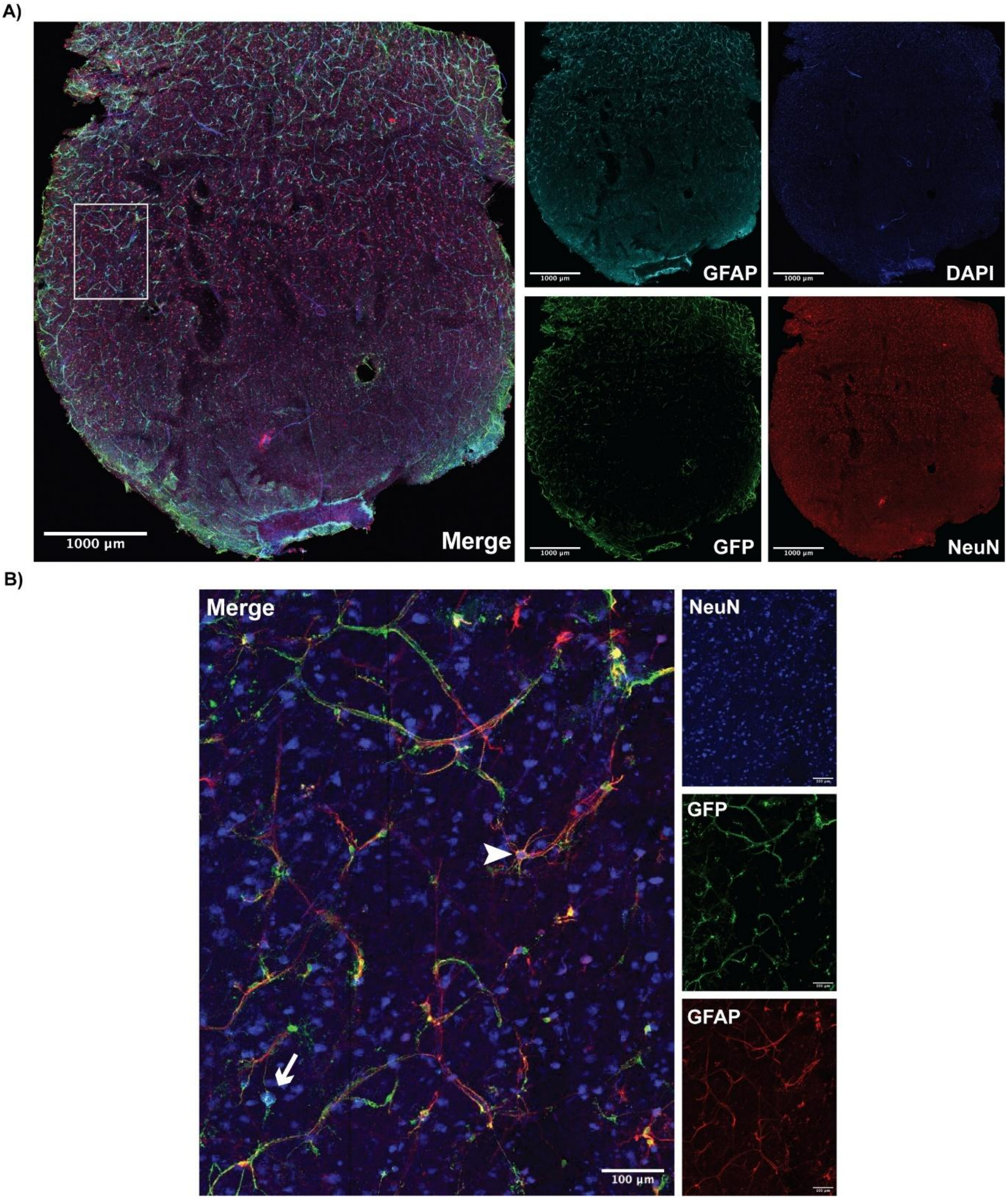


**Supplemental Figure 4: GFP Expression in human organotypic slices transduced with ShAAV.**

**A)** Representative confocal scanning images of human adult cortical slice transduced with AAVDJ-U6-CDKL5shRNA-CMV-eGFP (CDKL5 shRNA) and stained with GFAP (cyan),

GFP (green), NeuN (red) and the nuclear marker DAPI (blue). Scale bars represent 500 μm. Images were taken using a Zeiss LSM800 Airyscan confocal microscope.

**B)** Zoomed in area of A), position indicated by white box in A). In this image, DAPI is removed for easier visualisation: NeuN is blue, GFP is still green and GFAP is red. Scale bar represents 100 μm. Arrow shows GFP colocalization with NeuN (cyan, neuronal transduction) and pointed triangle shows GFP colocalization with GFAP (yellow, astrocytic transduction).
